# White matter hyperintensity genetic risk factor *TRIM47* regulates autophagy in brain endothelial cells

**DOI:** 10.1101/2023.12.18.566359

**Authors:** Sunny Hoi-Sang Yeung, Ralph Hon-Sun Lee, Gerald Wai-Yeung Cheng, Iris Wai-Ting Ma, Julia Kofler, Candice Kent, Fulin Ma, Karl Herrup, Myriam Fornage, Ken Arai, Kai-Hei Tse

## Abstract

White matter hyperintensity (WMH) is strongly correlated with age-related dementia and hypertension, but its pathogenesis remains obscure. GWAS identified TRIM47 at 17q25 locus as a top genetic risk factor for WMH formation. TRIM family is a class of E3 ubiquitin ligase with pivotal functions in autophagy, which is critical for brain endothelial cell (ECs) remodeling during hypertension. We hypothesize that TRIM47 regulates autophagy and its loss-of-function disturbs cerebrovasculature. Based on transcriptomics and immunohistochemistry, TRIM47 is found selectively expressed by brain ECs in human and mouse, and its transcription is upregulated by artificially-induced autophagy while downregulated in hypertension-like conditions. Using *in silico* simulation, immunocytochemistry and super-resolution microscopy, we identified the highly conserved binding site between TRIM47 and the LIR (LC3-interacting region) motif of LC3B. Importantly, pharmacological autophagy induction increased Trim47 expression on mouse ECs (b.End3) culture, while silencing *Trim47* significantly increased autophagy with ULK1 phosphorylation induction, transcription and vacuole formation. Together, we confirm that TRIM47 is an endogenous inhibitor of autophagy in brain ECs, and such TRIM47-mediated regulation connects genetic and physiological risk factors for WMH formation but warrants further investigation.

**SUMMARY STATEMENT:** TRIM47, top genetic risk factor for white matter hyperintensity formation, is a negative regulator of autophagy in brain endothelial cells and implicates a novel cellular mechanism for age-related cerebrovascular changes.

## INTRODUCTION

White matter hyperintensities (WMH) are abnormally strong signals detectable by magnetic resonance imaging (MRI) commonly found in the aging brain. The intense signals of WMH are the likely consequence of cerebrovascular changes with freely diffusible fluid originated from a leaky blood brain barrier (BBB), microbleeds (Alber et al., 2019; Roseborough et al., 2023). The age-related WMH burden increases with age and is significantly associated with cognitive decline (Au et al., 2006; Brown et al., 2021; Habes et al., 2016; Prins and Scheltens, 2015; Vergoossen et al., 2021). The radio-pathological correlation studies of WMH regions revealed a range of neuropathology including microinfarcts, astrogliosis, microglia activation, arteriolosclerosis and myelin loss (Fazekas et al., 1993; Gouw et al., 2008; Kalaria, 2016; Shim et al., 2015). The most consistent is BBB leakage as evident by the detection of fibrinogen in the penumbra region (Bridges et al., 2014; Hainsworth et al., 2017). At the WMH site, the cerebral capillary dysfunction triggers blood flow reduction, BBB disruption followed by fibrinogen breaching which ultimately causes oligodendrocyte loss (Montagne et al., 2018). While the gradual myelin loss would lead to cognitive impairment in age-related dementia (Tse and Herrup, 2017), the initial mechanism triggering the cerebrovascular changes underlying the WMH formation remains elusive.

Hypertension is one of the most prevalent conditions in the aging population, and it is the major risk factor for WMH formation (Chesebro et al., 2020; Hajjar et al., 2011; Verhaaren et al., 2013). The chronic hardening and narrowing of blood vessels in the cerebrovasculature can drive WMH-associated cognitive impairment even in young and asymptomatic populations (Hasan-Olive et al., 2019; Maillard et al., 2012). Cerebrovasculature is a highly dynamic system, whose flexibility is conferred by the constituent endothelial cells (ECs) (Kim et al., 2018). Together with pericytes and astrocytes, ECs are one of the three cellular constituents of BBB, which forms the interface between blood and brain (Zhao et al., 2015). At the vascular lumen, ECs constantly encounter shear stress of blood flow and adapt to hemodynamics by remodeling themselves through autophagy (De Meyer et al., 2015; Nussenzweig et al., 2015). In hypertensive conditions, the vascular tone and blood flow in the cerebrovasculature are impaired (Pires et al., 2013), and likely lead to autophagy failure in ECs (de Montgolfier et al., 2019; Vion et al., 2017).. As the restoration of autophagy was shown to reduce hypertension associated cerebrovascular dysfunction (Forte et al., 2020), we propose that impaired autophagy in ECs may be the underlying cause of hypertension-driven BBB leakage and WMH formation.

Despite the strong link between hypertension, autophagy failure and BBB leakage, not every hypertensive patient develops WMH (Hajjar et al., 2011). Population genetics fill this gap of knowledge. Independent GWAS (Genome wide association studies) identified SNPs (single nucleotide polymorphism) that confer significant risk to WMH development in the healthy aging population at locus at 17q25 (Fornage et al., 2011; Persyn et al., 2020; Sargurupremraj et al., 2020; Sarnowski et al., 2018; Traylor et al., 2019; Verhaaren et al., 2015). Among the six genes (*WBP2, TRIM65, TRIM47, MRPL38, FBF1,* and *ACOX1*) found at this locus, two are members of tripartite motif (TRIM) family. Importantly, a recent study combining GWAS and whole exome sequencing mapped to a missense variant of *TRIM47* that may cause the damaging effects towards WMH formation (Mishra et al., 2022). TRIM is a large family of E3 ubiquitin ligases with more than 80 members in the human genome. Members of TRIM family regulate protein turnover, quality control and degradation through ubiquitin-proteasome system during development, immunity and carcinogenesis (Hatakeyama, 2017). Surprisingly, TRIM family members are also heavily involved in the regulation of autophagy.

Many TRIM members were experimentally verified to interact with key autophagic adaptors including LC3, SQSTM1/p62 or ULK1/Beclin. Together, the proteins form a TRIM-containing complex during autophagosome formation for macroautophagy (Mandell et al., 2014), as well as selective microautophagy to precisely degrade intracellular organelles (Kimura et al., 2016). For example, TRIM65 is known to facilitate autophagy by controlling Atg7 expression in cancer cells (Pan et al., 2019). Yet, the role of TRIM47 in autophagy remains unknown, especially in endothelium. With the strong population genetic evidence, we hypothesized that the dysfunction of TRIM47 may cause autophagy failure in brain ECs, followed by BBB disintegration and WMH formation. In this study, we aim to characterize the role of TRIM47 in EC autophagy using a series of bioinformatic analysis coupled with experiments *in silico* and *in vitro*.

## METHODS

### Query of gene expression omnibus repository and scRNA-seq

The Gene Expression Omnibus (GEO) repository of NCBI was queried for datasets targeting hypertension, autophagy and/or endothelial cells. Four datasets (GSE108384, GSE199709, GSE180169 and GSE131712) were identified to test our hypothesis that *TRIM47* expression is altered in endothelial cells in hypertensive environments. The exploration and re-analysis were performed on the GEO2R R-based web application (Barrett et al., 2013) and GREIN:GEO RNA-seq Experiments Interactive Navigator (Mahi et al., 2019). The differential gene expression signature obtained in the pairwise comparison was identified. The significantly upregulated or downregulated genes reaching a false discovery rate or an adjusted P-value < 0.05 were then curated for gene ontology (GO) enrichment analysis by PANTHER-based platform (Mi et al., 2019).

### In silico analysis of TRIM47

To screen for the LC3-interacting region (LIR) motif among, the protein sequences of TRIM family members were queried in iLIR database, a web-based search engine LIR motif-containing proteins in eukaryotes (Jacomin et al., 2016; Kalvari et al., 2014). To simulate the three dimensional structure, the protein sequence of TRIM47 was mapped using Phyre2 server as described (Kelley et al., 2015). The best fit model generated by Phyre2, as well as the human and mouse TRIM47 protein model predicted by Alphafold artificial intelligence algorithm (Jumper et al., 2021) were docked with the human (PDB: 3VTU) or mouse LC3B (Alphafold: Q8C0E3) models using ClusPro web server 2.0 (Kozakov et al., 2017). The computed binding affinity between the protein structures were sorted based on scores measured by the estimated free energy coefficient (*E_balanced_*) and the binding cluster size. The best ranked protein-protein interaction structures were then explored, annotated and analyzed for contact area using software ICM Browser Pro (Molsoft L.L.C, (Abagyan et al., 1994)).

### Postmortem human brain tissue

A cohort of formalin fixed paraffin embedded (FFPE) postmortem brain tissues were used were kindly provided by the Neuropathology Core of Alzheimer’s Disease Research Center at University of Pittsburgh Medical Center with approvals from the Committee for Oversight of Research and Clinical Training Involving Decedents at University of Pittsburgh. The postmortem tissues were of frontal cortex origin (Brodmann area 9) from two males and seven females of Alzheimer’s disease (AD, Braak Stage VI) and age-matched controls (Braak Stage 0-III) between the ages of 73 and 90 years old. The postmortem interval was between 3.5 and 19 hours. All tissues were sectioned at 12 µm and stored at room temperature until use.

### Animal study

The animal experiment was approved by the Animal Subjects Ethics Sub-Committee at The Hong Kong Polytechnic University (PolyU) (ASESC Case No.: 20-21/167-HTI-R-GRF) with a valid license from the Department of Health in Hong Kong. Wildtype mice (WT, C57BL/6J) were housed in individually ventilated cages (Techniplast, Italy) in a temperature and humidity-controlled environment with food (PicoLab Diet 5053, LabDiet Inc, MO, USA) and water *ad libitum* on a 12-hour light/dark cycle The brain tissue was extracted from WT mice at ten months of age (n = 3). Briefly, animals were deeply anesthetized prior to transcardial perfusion with phosphate buffer saline, as described earlier (Mok et al., 2023). The fresh brain was fixed by paraformaldehyde (4%, 24 hours) and then cryoprotected in PBS-sucrose (30% w/v, 72 hours) at 4 °C before embedded for cryosectioning on SuperFrost-plus slides at 10 μm thickness.

### Immunohistochemistry-Immunofluorescence

For human tissue, the paraffin-fixed sections were dewaxed with xylene before rehydration in a series of ethanol/phosphate-buffered saline. The rehydrated sections were antigen retrieved with citrate buffer (pH 6.0) at 95°C for 10 min. After cooling and rinsing, the non-specific antibody binding was then blocked using 10% donkey serum at room temperature for one hour, followed by overnight incubation at 4°C with the primary antibody against TRIM47 (1:100, rabbit polyclonal, ab72234, Abcam). The tissue sections were washed and then incubated in AlexaFluor-conjugated secondary antibody for 1 h at room temperature with nuclear counterstained by PI. All immunofluorescence image of the human tissues was captured in University of Pittsburgh using a Nikon A1R confocal system based on an Eclipse Ti2 microscope equipped by a 20x (Nikon N Plan Apo 20x/0.75), 40x (Nikon N Plan Apo 40x/0.95) dry objectives, or 60× (1.4 Numerical Aperture) oil objective lens and a camera (at 512 µm × 512 µm) operated by NIS-Elements AR software. Three Region-Of-Interests were captured per specimen for quantification.

For mouse tissue, the frozen sections were washed and blocked with normal donkey serum (10% v/v) in phosphate buffer saline (PBS) with 0.3% Triton X-100 for 1 hour at room temperature. Then blocked tissues were then incubated with specific primary antibodies against anti-CD31 antibody [P2B1] ab24590 (1:500, mouse monoclonal, ab72234, Abcam) and anti-TRIM47 (1:500, rabbit polyclonal, ab72234, Abcam) overnight at 4 °C. The sections were then washed with PBS before incubation with incubated secondary antibodies against mouse or rabbit IgG conjugated with Alexa Fluor 488 or 555 or 647 (Thermo Fisher Scientific, MA) for one hour at room temperature. All nuclei were counterstained with DAPI before mounting in Hydromount (National Diagnostics, GA, USA) with coverslips. Whole brain tissue sections were taken by a Nikon Eclipse Ti2-E Live-cell Fluorescence Imaging System at University Research Facility in Life Sciences as described below. The image was analyzed by ImageJ (FIJI 1.53, National Institute of Health, MD, USA).

### Cell culture study

To study the function of Trim47 in autophagy, an endothelial cell line derived from mouse brain, bEnd.3 (CCL-2299, ATCC), was used (Montesano et al., 1990). bEnd.3 cells were maintained in complete Dulbecco’s Modified Eagle Medium supplemented with 10% (v/v) fetal bovine serum and 1% (v/v) and penicillin/streptomycin mixture in incubator at 37°C with 5% CO_2_. To passage, the cells were detached with 0.25% (v/v) trypsin before replating. To study autophagy, bEnd.3 cells were treated with either 3-methyladenine (3-MA, #1292 Tocris, MN, USA) or rapamycin (Rapa, #3977 Tocris, MN, USA) for 24 hours, or gene silencing described below, before downstream analysis.

### Gene silencing

To study the loss-of-function, *Trim47* expression in bEnd.3 cells was silenced by SMARTpool siRNA (siGENOME Mouse Trim47, Gene ID 217333, M-056879-01-0020) or control siRNA (siGENOME non-targeting, D-001206-13-20) using DharmaFECT 1 Transfection Reagent (T-2001-02, Horizon Discovery, USA) according to manufacturer’s instruction.

### Western blotting

To obtain cell lysates, the cells were lysed by in radioimmunoprecipitation assay buffer (20-188, Millipore, CA, USA) with protease and phosphatase inhibitors (Roche, Hong Kong). After quantification, the protein samples were normalized (Bio-Rad Protein Assay, 5000001, Bio-Rad, CA, USA) and 30 µg were loaded into a 7.5 to 12% sodium dodecyl sulfate (SDS)-polyacrylamide gel for electrophoresis. For the routine immunoblotting, the protein was heated for 5 minutes at 95°C for denaturation. For non-denaturing immunoblotting, the heating step was omitted and the SDS was omitted from the sample buffer. The resolved proteins were transferred to a semi-dry polyvinylidene difluoride (PVDF) membrane using Bio-Rad Trans-Blot Turbo Transfer System (1704150, Bio-Rad, CA, USA). After blocking non-specific bindings with 5% non-fat milk, the blot was probed against TRIM47 (1:500, rabbit polyclonal, ab72234, Abcam), LC3B (1:1000, E5Q2K, mouse monoclonal #83506, Cell signaling technology/CST), ULK1 (1:1000, D8H5, rabbit monoclonal #8054S, CST), p-ULK (Ser555) (1:1000, D1H4, rabbit monoclonal, #5869, CST), p62 (1:1000, D1Q5S, rabbit monoclonal, #39749, CST), p-p62 (Ser349) (1:1000, E7M1A, rabbit monoclonal, #16177, CST) and GABARAP (1:1000, E1J4E, rabbit monoclonal, #13733, CST). After overnight incubation, the probed blots were incubated with horseradish peroxidase-conjugated secondary immunoglobulins (Cell Signaling Technology, MA, USA). After washes with tris-buffer saline-Tween, the protein signals were captured by chemiluminescent substrates (SuperSignal™ West Pico or Extended Dura or Femto Maximum Sensitivity Substrate, Thermo Fisher Scientific, MA, USA) and the blot image was acquired using Bio-Rad ChemiDoc MP Imaging System (12003154, Bio-Rad, CA, USA) with a continuous exposure for 5 minutes. To quantify the protein expression level, the density and the size in kilodalton (kDa) of each individual band was measured by ImageJ (FIJI, v1.53, National Institute of Health, MD, USA).

### Gene expression

The total RNA was isolated from the cells using RNeasy Mini Kit (Qiagen, Germany, 74106). Any remaining genomic DNA in the RNA sample was digested by DNase I (1 Unit /μL, Thermo Fisher Scientific, MA, USA). To generate cDNA, reverse transcription was performed with High-capacity RNA-to-cDNA kit (Applied Biosystem, Thermo Fisher Scientific, MA, USA). The genes of interest and the housekeeping gene, *Gapdh*, was amplified and analysed in real-time polymerase chain reaction on a Roche Light Cycler 480 system with TB Green® Premix Ex Taq™ II (TaKaRa Bio Inc., Japan). The specific primers were predesigned and validated by PrimerBank of Massachusetts General Hospital (Spandidos et al., 2010), with the primer sequences detailed in Table S1.

### Immunocytochemistry

For microscopic examination, immunocytochemistry was performed as described (Mok et al., 2023). The bEnd.3 cells plated on coverslips were fixed by 4% paraformaldehyde (pH 7.2) for 30 minutes. After washing with PBS, the potential non-specific binding was blocked with 5% (v/v) normal donkey serum (D9663, Sigma-Aldrich) in PBS containing 0.3% Triton X-100 for 30 minutes at room temperature. The specimen was then incubated with primary antibody against LC3B (83506S, 1:400, Cell Signaling Technology) as well as TRIM-47 (ab72234, 1:400, Abcam), p-p62 Ser349 (1:800, CST, #16177S), p-ULK1 Ser555 (1:250, CST, #5869) or GABARAP (1:200, CST, #13733) for two hours, followed by fluorescent visualization with donkey anti-mouse or anti-rabbit Alexa fluor-488 or -555 (A31572, A21202, Thermo Fisher Scientific, MA, USA). The stained coverslip was further incubated with DAPI (4’,6-diamidino-2-phenylindole, 1 µg/mL) for 5 minutes before mounted on glass slide with Hydromount aqueous mounting medium (HS106100ML, National Diagnostics). The coverslips were examined with a Nikon Eclipse Ti2-E Live-cell Fluorescence Imaging System at University Research Facility in Life Sciences. The system is equipped with DS-Qi2 monochrome camera and a multi-wavelength LED illumination system (CoolLED pE-4000) with blue (Ex: 380/55), green (Ex: 470/30), red (Ex: 557/35) and far-red (Ex: 620/60) filters with a Nikon Plan Apo λ 20x Objective (N.A. 0.75, R.I. 1.0). Duplicate coverslips were included for each condition and a total of five images (2424 x 2424 pixel) from random fields (882.5 µm x 882.5 µm) was acquired using Nikon NIS elements software, (V5.21.03 Nikon Instruments, NY, USA). Image analysis was performed by QuPath (Bankhead et al., 2017) and Cellprofiler (McQuin et al., 2018).

### Quantification of autophagic puncta

The quantification of ATG8 puncta is a standard approach to measure the level of autophagic flux (Klionsky et al., 2021). To quantify autophagy, the immunocytochemistry image was analyzed by the open-source CellProfiler Software of version 4.2.5 (McQuin et al., 2018). Briefly, the triple immunofluorescent images (2424×2424 pixels at 0.36 µm/pixel) were separated into three channels and converted into greyscale. The primary objects of the nucleus, green or red puncta were identified and related in each cell. The related features were then classified, filtered and masked for the measurement of signal colocalization. In each masked cell, the second and tertiary objects segmenting the cytoplasm and the cytoplasmic puncta were then identified and classified, with the number of cytoplasmic puncta in each positive cell quantified for analysis.

### Structured Illumination Microscopy imaging

Super resolution microscopy was performed using a Structured Illumination Microscopy (SIM) system at University Research Facility in Life Sciences. The Nikon N-SIM was built on a fully motorized Eclipse Ti inverted microscope body equipped with Nikon LU-NV MultiLaser (TIRF), ORCA-Flash4.0 V2+ sCMOS camera and CFI SR HP Plan Apo 100× (NA = 1.35) oil objective lens. The XY resolution is 115 nm and Z resolution is 250 nm in SIM mode (Nikon Instruments, NY, USA). To prepare the samples for N-SIM, cells were plated on 1.5H 13 mm Marienfeld no. round coverglass as described above but with the specimen mounted using ProLong Diamond antifade mountant (Thermo Fisher Scientific, MA, USA). The acquired datasets with an array of 1024 × 1024 pixels across the Z steps of 1 micron were computationally reconstructed using built-in algorithm in NIS-Elements AR software.

### Statistics

For each experiment, at least three independent biological replicates were included, and the total number of experiments is denoted as *n* number in each graph. A Grubbs’ test was performed to identify outliners identification. For comparisons of one variable between two groups, an unpaired t-test was performed. For comparisons of one variable in more than two groups, a one-way analysis of variance (ANOVA) was performed with a post-hoc tests for multiple comparison. For comparisons of two or more variables in two groups or more, a two-way ANOVA with a post-hoc multiple comparison was conducted. All statistical analyses were performed using GraphPad Prism software version 10.0 (GraphPad Software Inc. MA, USA). All statistical analyses were two-tailed with, and the significance level was set at p < 0.05. The details of the analysis and post-hoc tests are stated in each figure legend.

## RESULTS

To test our hypothesis that TRIM47 dysfunction in the brain endothelium leads to WMH formation, we first sought to confirm if *TRIM47* was selectively expressed in brain EC. We queried multiple single cell RNAseq (scRNAseq) database of human and mouse brain tissues using the single cell portal (Fig. 1). In two independent scRNAseq libraries of human prefrontal cortices from healthy adult subjects (Habib et al., 2017; Russell et al., 2023), *TRIM47* expression was highly enriched in the brain ECs and astrocytes and essentially absent in neurons and other brain cell types (Fig. 1A). Further exploration of an additional scRNAseq library of normal human blood brain barriers further showed that brain ECs from arterioles, capillaries and veins similarly express TRIM47 and their level is stronger than that of pericytes and fibroblast (Fig. 1B, C, (Garcia et al., 2022)), while the level in smooth muscle cells was highly variable. Using immunohistochemistry on a cohort of healthy and AD human cerebral cortex, we confirmed that TRIM47 signals are detectable in the endothelium of the capillary and arteriole-like structures (Fig.1D), but not in neuron or neurophil. To confirm *Trim47* expression pattern in mouse brain, we explored the scRNAseq library of mouse whole brain (C57BL/6J, 2-3 months of age, n = 8, (Ximerakis et al., 2019)) and a scRNAseq library of mouse whole cerebral cortex (C57BL/6J, 1 month of age, n = 1, (Ding et al., 2020)). We found that *Trim47* is strongly expressed by mouse brain ECs in these independent libraries (Fig. 1E). This is confirmed by an additional query on a third library of scRNAseq of transcriptions from nine adult mouse brain regions (i.e. frontal cortex, posterior cortex, substantia nigra, striatum, thalamus, globus pallidus, entopeduncular, hippocampus and cerebellum; C57Blk6/N, 2 months of age). In each brain region, the highest level of *Trim47* transcription was found in the Fit1^+^ ECs population (Fig. 1F, http://dropviz.org/ (Saunders et al., 2018)). In both datasets, pericytes also express Trim47 at a lower level. Using immunohistochemistry, we confirmed that Trim47 is selectively expressed by CD31^+^ brain endothelial cells in wildtype C57BL/6J mice at 10 months old (Fig. 1G). These observations from six independent scRNAseq datasets agree with our histology experiments and suggest that TRIM47 is selectively expressed by brain ECs of human and mouse.

**Figure 1.**
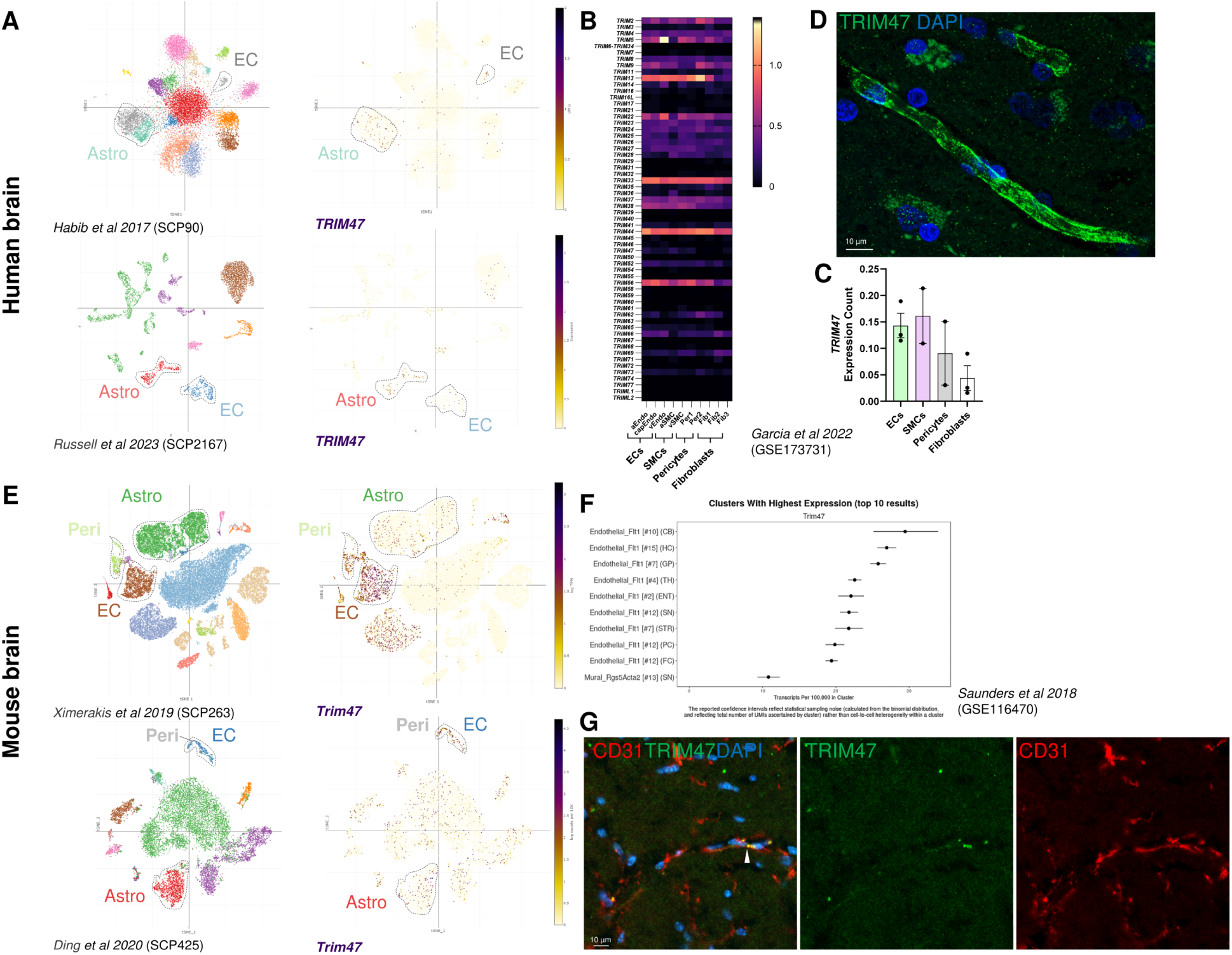
*TRIM47* expression is enriched in brain ECs in human and mouse. **A** Single cell portal-based scRNAseq exploration with t-SNE (t-distributed stochastic neighbour embedding) visualization of gene expression in human brain tissue (accession: SCP90, SCP2167). *Left*, t-SNE representation of all detect brain cell types *Right*, t-SNE representation of human *TRIM47* expression enriched in the brain endothelial cells and astrocytes. All cell labels are colour coded to the tSNE map. **B** Exploration of *TRIM* family transcription in a scRNAseq library of healthy human cerebrovascular cells (GSE173731), with *TRIM47* expression shown on **C**. **D** Representative *TRIM47* (green) immunohistochemistry image of the healthy human frontal cortex confirming the specific expression on endothelial cells. **E** Single cell portal-based scRNAseq exploration with t-SNE visualization of gene expression in mouse (wildtype) brain tissues (accession: SCP263, SCP425). *Left*, t-SNE representation of all detected brain cell types *Right*, t-SNE representation of mouse *Trim47*expression enriched in the Fit1^+^ endothelial cells. **F** Exploration of in a third scRNAseq of a mouse (wildtype) brain across nine regions showing brain ECs has the highest *Trim47* expression in all regions. **G** Representative immunohistochemistry image of the wildtype mouse corpus callosum confirming the specific Trim47 expression (green) on CD31^+^ endothelial cells (red). aEndo, arterioles endothelial cells; capEndo, capillary endothelial cells; vEndo, venule endothelial cells; aSMC, arterioles smooth muscle cell; vSMC, venule smooth muscle cell; Per1/Per2, Pericytes; Fib1/Fib2/Fib3, fibroblasts. FC, frontal cortex; PC, posterior cortex; SN, substantia nigra; STR, striatum; TH, thalamus; GP, globus pallidus; ENT, entopeduncular; HC, hippocampus and CB, cerebellum.

Recall that ECs require autophagy to remodel the vascular structure upon stimulation by luminal shear stress (De Meyer et al., 2015; Nussenzweig et al., 2015). We then characterized *TRIM47* expression in ECs during autophagy and autophagy-inducing conditions through a meta-analysis of TRIM family members in independent RNA-seq datasets. First, in adenovirus-mediated TFEB (transcription factor EB, the master regulator of autophagy) overexpression in primary human EC culture (HUVEC, human umbilical vein endothelial cells), autophagy was significantly induced with increased EC migration and tube formation (Fan et al., 2018). The meta-analysis of the microarray dataset (GSE108384) showed that 27 *TRIM* family members were significantly and differentially expressed in these autophagic human ECs (adjusted P value < 0.05). Among the differentially expressed *TRIM* members, *TRIM47* was the third most upregulated gene (Log_2_ fold change = 1.16, Fig. 2A). To test our hypothesis that TRIM47 function may be associated with such blood flow-related autophagy, we inquired *TRIM* family members expression in a human aortic ECs model with or without luminal shear stress simulation (12 dyn/cm^2^, 12 hours; RNA-seq, GSE199709) (Meng et al., 2022). In gene ontology analysis, macroautophagy (GO:0034262) was significantly enriched by the differentially upregulated genes expressed by ECs immediately after 12 hours shear stress treatment. In this autophagic condition, 14 out of the 49 detectable *TRIM* family members were significantly and differentially expressed by the ECs, where *TRIM47* was the sixth most significantly downregulated family member (Log_2_ fold change = -0.41, Fig. 2B), but such reduction was abolished after 24 hours of recovery in static medium. The details of these re-analyses can be found in Supplementary Tables S2 and S3.

**Figure 2.**
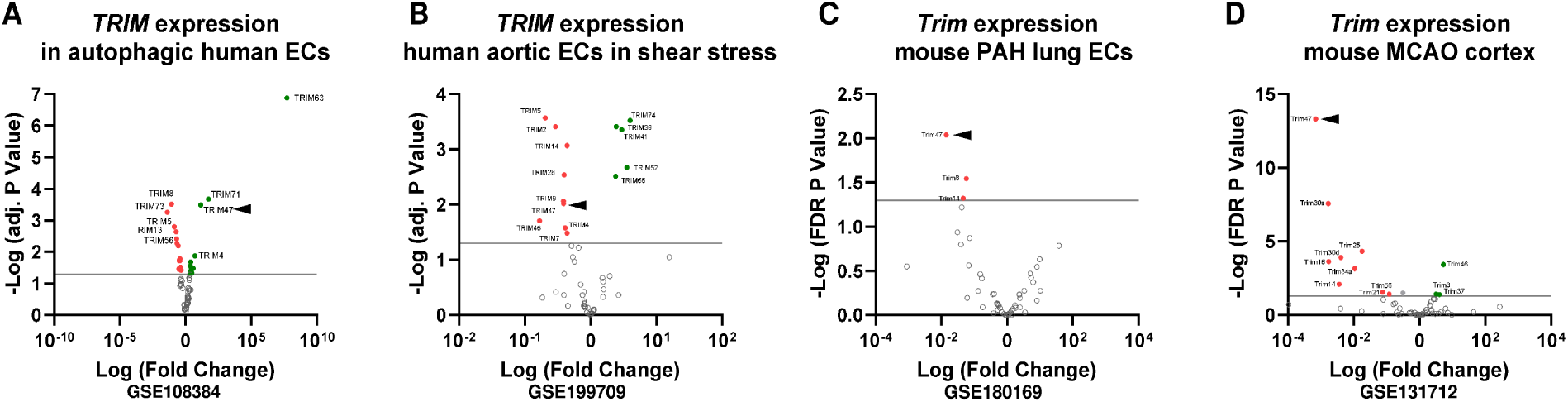
TRIM family transcription across hypertensive and cardiovascular disorder models in human and mouse. **A** *TRIM47* expression was significantly regulated in primary HUVEC cells upon overexpression of master autophagy transcription factor TFEB **B** *TRIM47* expression was downregulated in primary human aortic ECs upon laminar shear stress stimulation, but the level return to normal after 24 hour of recovery **C** A significant downregulation of *Trim47* expression was found in mouse lung endothelial cells after 3 weeks of pulmonary arterial hypertension modelling **D** A significant downregulation of *Trim47* expression was found in mouse middle cerebral artery occlusion modelling 24 hours upon injury, but the level return to normal after 28 days of recovery. Extended data are presented see Table S2 and Table S3.

To confirm if the association between *TRIM47* and autophagy is conserved in the mouse model, we investigated *Trim47* expression in a pulmonary hypertension model (PAH, (Rodor et al., 2022)). The lung ECs in the PAH model were sorted by flow cytometry, and we identified a significant enrichment of the autophagic process in the hypertensive condition, as in humans (Fig. 4C and Table S3). Among the 49 detectable *Trim* family members, three were significantly downregulated in the ECs and *Trim47* ranked number one (Log_2_ fold change = 1.85, RNA-seq, GSE180169). As changes in blood flow of the cerebrovasculature will cause autophagy failure in brain ECs (Pires et al., 2013; Vion et al., 2017), we inquired if *Trim47* expression is altered in an ischemic brain injury model. In a mouse model of transient middle cerebral artery occlusion (MCAO, C57BL/6J, 8–10 weeks of age, occlusion for 1 hour, (Wu et al., 2019)), 14 out of the 59 detectable *Trim* family members were significantly and differentially expressed in the ipsilateral cerebral cortex 24 hours post-injury, where *Trim47* was the most significantly downregulated member (Log_2_ fold change = -3.163, RNA-seq, GSE131712). Intriguingly, such downregulation was abolished after 28 days of recovery in the ipsilateral cortex. In a similar mouse MCAO model (129S6/SvEv, 8-10 weeks of age, occlusion for 30 minutes, (Wegner et al., 2020)), the brain ECs isolated from the injured brain after 72 hours of recovery similarly showed no changes in any *Trim* family members and no autophagic process was significantly enriched among the differentially expressed genes (FDR-based, RNA-seq, GSE122345, Table S2 and S3). Together, these independent datasets suggested the TRIM47 transcription in ECs is significantly upregulated when autophagy is artificially induced, but significantly downregulated when the hypertension- or cerebrovascular dysfunction-related autophagy begins.

The bioinformatic inquiry above suggested that TRIM47 belongs to the group of TRIM family members not only involved in ubiquitination-mediated degradation, but also in autophagy. Indeed, many of TRIM members interacts with human ATG8 (LC3), p62/SQSTM1 and/or ULK1/Beclin1 (TRIM -1, -5, -6, -13, -16, -17, -20, -21, -22, -28, -33, -40, -41, -45, -49, -50, -55, -61, -63 and -76) (Hatakeyama, 2017; Mandell et al., 2014) and forms a TRIM-containing complex during autophagosome formation. Nevertheless, TRIM47 was not detected as one of these autophagic regulating members. To clarify, we performed a systemic screening of human and mouse TRIM protein sequences for the LIR motifs using iLIR database (Table 1) (Jacomin et al., 2016; Kalvari et al., 2014). LIR motif is the consensus amino acid sequence responsible for autophagic functions. This consensus sequence includes WxxL and xLIR, where **<u>W</u>**xx**<u>L</u>** is the canonical consensus sequence of LC3B interacting site where x can be any amino acid among the four residues. The consensus was then extended as xLIR, where the six amino acid residues can be any combination of (ADEFGLPRSK)(DEGMSTV)(**<u>WFY</u>**)(DEILQTV)(ADEFHIKLMPSTV)(**<u>ILV</u>**). Such motif would also have a higher probability to bind to LC3 if it is located within an anchor, a disordered region with high potential to be stabilized upon binding (Kalvari et al., 2014). Using the position specific scoring matrix (PSSM) as the measurement, we found that human TRIM47 attained a PSSM score of 14 and it is within the threshold range of the verified xLIR molecules (PSSM = 13-17) (Kalvari et al., 2014). Importantly, the xLIR (NQWEQL) motif in human TRIM47 fit within an anchor region. Such potential LC3 interacting site was conserved in mouse Trim47 (SQWEQL). Although this motif is not embedded within an anchor, the xLIR in Trim47 attained a PSSM score of 16 and within the threshold of verified xLIR containing proteins. Intriguingly, a low PSSM score was found in human TRIM65 (9) and mouse Trim65 (8), the other TRIM family member identified in the 17q25 locus.

**Table 1.** Screening of LIR (LC3-interacting region)-motif among known autophagy-associated TRIM family members in human and mouse. PSSM: position-specific scoring matrix score; LIR = WxxL or xLIR; **W**xx**L**, LIR consensus sequence where x can be any amino acid among the four residues; xLIR, LIR consensus sequence where the six residues can be any amino acids as follow (ADEFGLPRSK)(DEGMSTV)(**WFY**)(DEILQTV)(ADEFHIKLMPSTV)(**ILV**). SQSTM encodes Sequestosome 1(or p62) which is a verified LC3 interacting substrate and serves as the positive control.

| Human |  |  |  |  |  |  |  | Mouse |  |  |  |  |  |  |  |
| --- | --- | --- | --- | --- | --- | --- | --- | --- | --- | --- | --- | --- | --- | --- | --- |
| Gene Symbol | NCBI Protein accession | Motif | Start | End | Pattern | PSSM Score | LIR in Anchor | Gene Symbol | NCBI Protein accession | Motif | Start | End | Pattern | PSSM Score | LIR in Anchor |
| *SQSTM1* | NP_003891.1 | xLIR | 336 | 341 | DDWTHL | 24 | Yes | *Sqstm1* | NP_035148.1 | xLIR | 338 | 343 | DDWTHL | 24 | Yes |
| *TRIM47* | NP_258411.2 | WxxL | 379 | 384 | NQWEQL | 14 | Yes | *Trim28* | EDL38082.1 | WxxL | 515 | 520 | EDYNLI | 13 | Yes |
| *TRIM28* | AAH04978.1 | WxxL | 515 | 520 | EDYNLI | 13 | Yes | *Trim63* | NP_001034137.2 | WxxL | 305 | 310 | MDYFTL | 11 | Yes |
| *TRIM63* | NP_115977.2 | WxxL | 305 | 310 | MDFFTL | 10 | Yes | *Trim20* | XP_006522424.1 | WxxL | 227 | 232 | EVYVYL | 8 | Yes |
| *TRIM41* | NP_291027.3 | WxxL | 315 | 320 | SEFGRL | 7 | Yes | *Trim41* | NP_663352.2 | WxxL | 315 | 320 | SEFGRL | 7 | Yes |
| *TRIM20* | NP_000234.1 | xLIR | 523 | 528 | SEWELL | 20 | No | *Trim21* | NP_001076021.1 | xLIR | 5 | 10 | KMWEEV | 18 | No |
| *TRIM13* | NP_998755.1 | WxxL | 284 | 289 | SQWEDI | 17 | No | *Trim47* | NP_001192010.1 | WxxL | 383 | 388 | SQWEQL | 16 | No |
| *TRIM21* | NP_003132.2 | WxxL | 9 | 14 | MMWEEV | 16 | No | *Trim25* | NP_033572.2 | xLIR | 302 | 307 | DEFEFL | 16 | No |
| *TRIM25* | NP_005073.2 | xLIR | 303 | 308 | DEFEFL | 16 | No | *Trim6* | NP_001013637.1 | WxxL | 298 | 303 | SYWVDV | 14 | No |
| *TRIM49* | NP_065091.1 | WxxL | 41 | 46 | LNWQDI | 15 | No | *Trim5* | NP_001297531.1 | WxxL | 296 | 301 | RYWVQV | 13 | No |
| *TRIM5* | NP_001397887.1 | WxxL | 183 | 188 | ADFEQL | 14 | No | *Trim50* | NP_839971.1 | xLIR | 177 | 182 | REFQEL | 13 | No |
| *TRIM6* | NP_001003818.1 | WxxL | 326 | 331 | SYWVDV | 14 | No | *Trim13* | NP_001157692.1 | WxxL | 284 | 289 | SQWGDI | 12 | No |
| *TRIM50* | NP_835226.2 | xLIR | 177 | 182 | REFQEL | 13 | No | *Trim17* | NP_112449.1 | xLIR | 187 | 192 | EEFQKV | 11 | No |
| *TRIM22* | NP_006065.2 | WxxL | 173 | 178 | KNYIQI | 9 | No | *Trim55* | NP_001074750.1 | WxxL | 213 | 218 | EKFDHL | 10 | No |
| *TRIM55* | NP_908973.1 | WxxL | 213 | 218 | EKFDYL | 9 | No | *Trim65* | NP_848917.2 | WxxL | 344 | 349 | NQYLHL | 8 | No |
| *TRIM65* | NP_775818.2 | xLIR | 333 | 338 | LTFDPV | 9 | No | *Trim40* | NP_001028407.1 | WxxL | 170 | 175 | LPWHWL | 7 | No |
| *TRIM17* | NP_001020111.1 | WxxL | 401 | 406 | STFSAL | 9 | No |  |  |  |  |  |  |  |  |
| *TRIM40* | NP_001273562.1 | WxxL | 238 | 243 | VIYPQL | 0 | No |  |  |  |  |  |  |  |  |

To confirm the putative autophagic function of TRIM47, we performed a structural analysis of the potential TRIM47-LC3B interactions *in silico*. As no prior X-ray crystallography-based or NMR-based study is available, we first predicted the protein folding structure of human TRIM47 using Phyre2 Protein Fold Recognition Server (Kelley et al., 2015). Under the intensive mode of simulation, the final TRIM47 model mapped on the templates of TRIM20 (PDB: 4CG4; Sequence identity 16%, Confidence 100%), TRIM25 (PDB: 6FLN; Sequence identity 24%, Confidence 100%) and TRIM28 (PDB: 6QAJ; Sequence identity 19%, Confidence 100%), see Fig 3A-C. This model was docked with human LC3B protein in ClusPro server (Kozakov et al., 2017), and we identified the potential interactions between the TRIM47 xLIR motif (N379 and Q380 of NQWEQL) and LC3B (K55 and R14, PDB: 3VTU) with a distance of 2.7 – 2.9 Å (Table S4). To validate this interaction, we repeated the LC3B docking experiment with a new artificial intelligence based human TRIM47 model predicted by Alphafold (Jumper et al., 2021) (Alphafold: Q96LD4. Fig. 3D-F). In this docking experiment, we similarly identified potential interactions between the TRIM47 xLIR motif and LC3B, but through different residues, where the W381, Q383 and L384 of NQWEQL bound to I21, R72 and V93 of LC3B (PDB: 3VTU) with a distance between 2.3 – 4.1 Å. Importantly, the putative NQWEQL interacting contacts at R14 and R72 are identical to the known LC3B binding site at hydrophobic pocket 1 and 2 (HP1 near helix α1, HP2 near helix α3; equivalent to residue R10 and R69 of PDB: 5WRD, respectively (Wirth et al., 2019)). Furthermore, the K55 contact site is also a known basic residue and electrostatic binding site important for p62/SQSTM1-LC3 interactions (equivalent to residue K50; (Johansen and Lamark, 2011)). To test if Trim47-LC3B interaction is conserved in mouse, we docked mouse Trim47 (Alphafold: Q8C0E3) and mouse LC3B (Alphafold: Q9CQV6) and identified a similar binding as in human (Fig 3G-I). Although the binding residues between the mouse xLIR in Trim47 (SQWEQL) and LC3 at F80 and M88 were not known sites of interaction important for autophagy, the potential interaction (Energy weighted score) is comparable to that in human (See Table S4). Together, this docking simulation confirms that TRIM47 interacts with LC3B in human and mouse.

**Figure 3.**
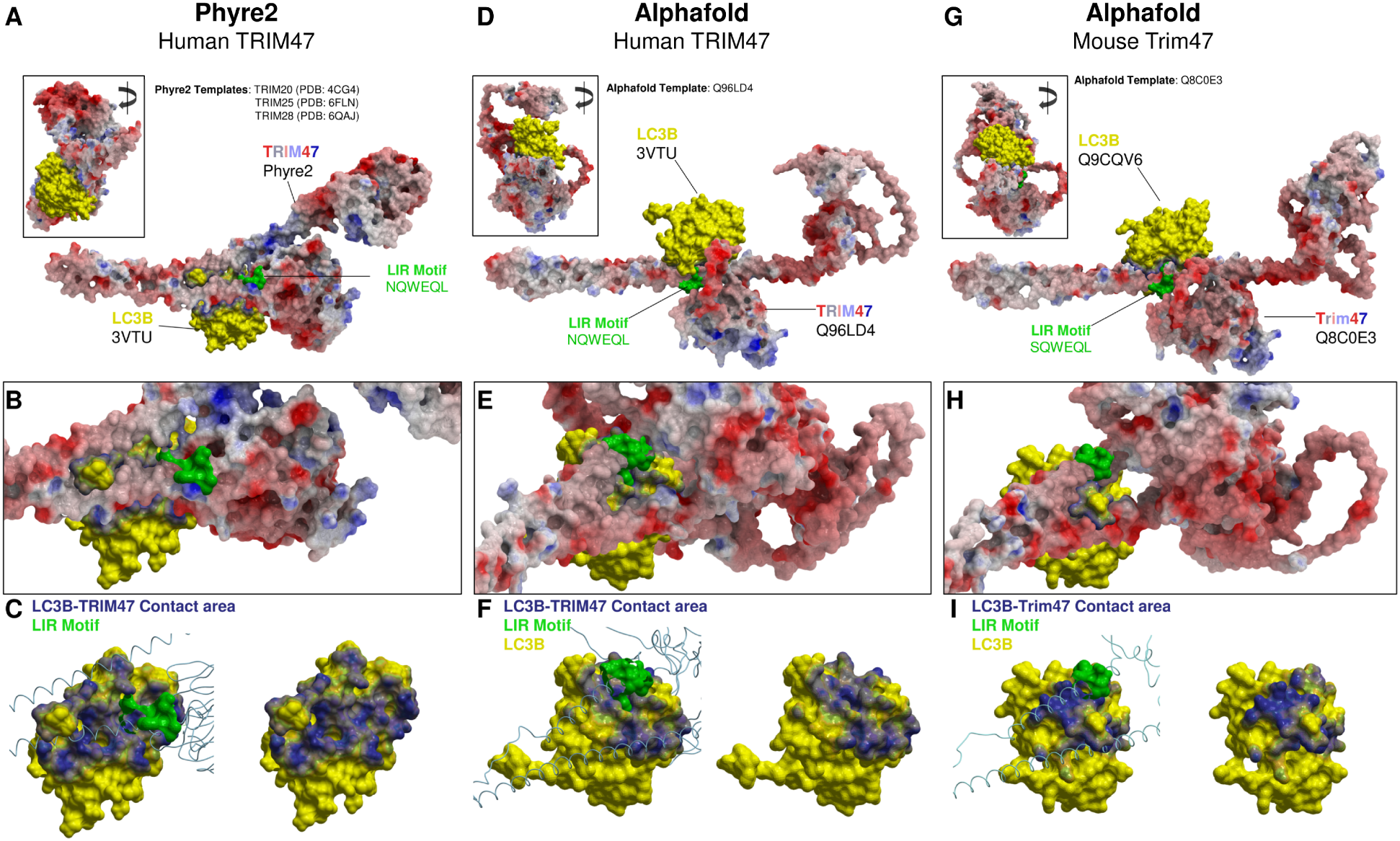
Simulation of xLIR motif-LCB interactions in TRIM47 *in silico*. **A** Phyre2 generated 3D model of TRIM47 (Red-grey-blue on electrostatic charge cartoon mesh; model based on PDB: 4CG4, 6FLN and 6QAJ) docked with human LC3B (Yellow; PDB: 3VTU) using ClusPro server (Kozakov et al., 2017).The xLIR sequence NQWEQL highlighted in green. Right 90° rotation in inset **B** Magnified view of the interaction site between xLIR and LC3B. **C** xLIR-LC3B interactions with (*left*) or without (*right*) the ribbon of xLIR (green), where the xLIR-LC3B contact area highlighted in blue. **D** Human TRIM47 generated by artificial intelligence (Red-grey-blue on electrostatic charge cartoon mesh; Alphafold: Q96LD4) docked with human LC3B (Yellow; PDB: 3VTU). The xLIR sequence NQWEQL is highlighted in green. Right 90° rotation in inset **E** Magnified view of the interaction site between xLIR and LC3B. **F** xLIR-LC3B interactions with (*left*) or without (*right*) the ribbon of xLIR (green), where the xLIR-LC3B contact area is highlighted in blue. **G** Alphafold-based Mouse Trim47 (Red-grey-blue on electrostatic charge cartoon mesh; Alphafold: Q9CQV6) docked with mouse LC3B (Yellow; Alphafold: Q8C0E3). The xLIR sequence SQWEQL highlighted in green. Right 90° rotation in inset **H** Magnified view of the interaction site between xLIR and LC3B. **I** xLIR-LC3B interactions with (*left*) or without (*right*) the ribbon of xLIR (green), where the xLIR-LC3B contact area is highlighted in blue.

**Figure 4.**
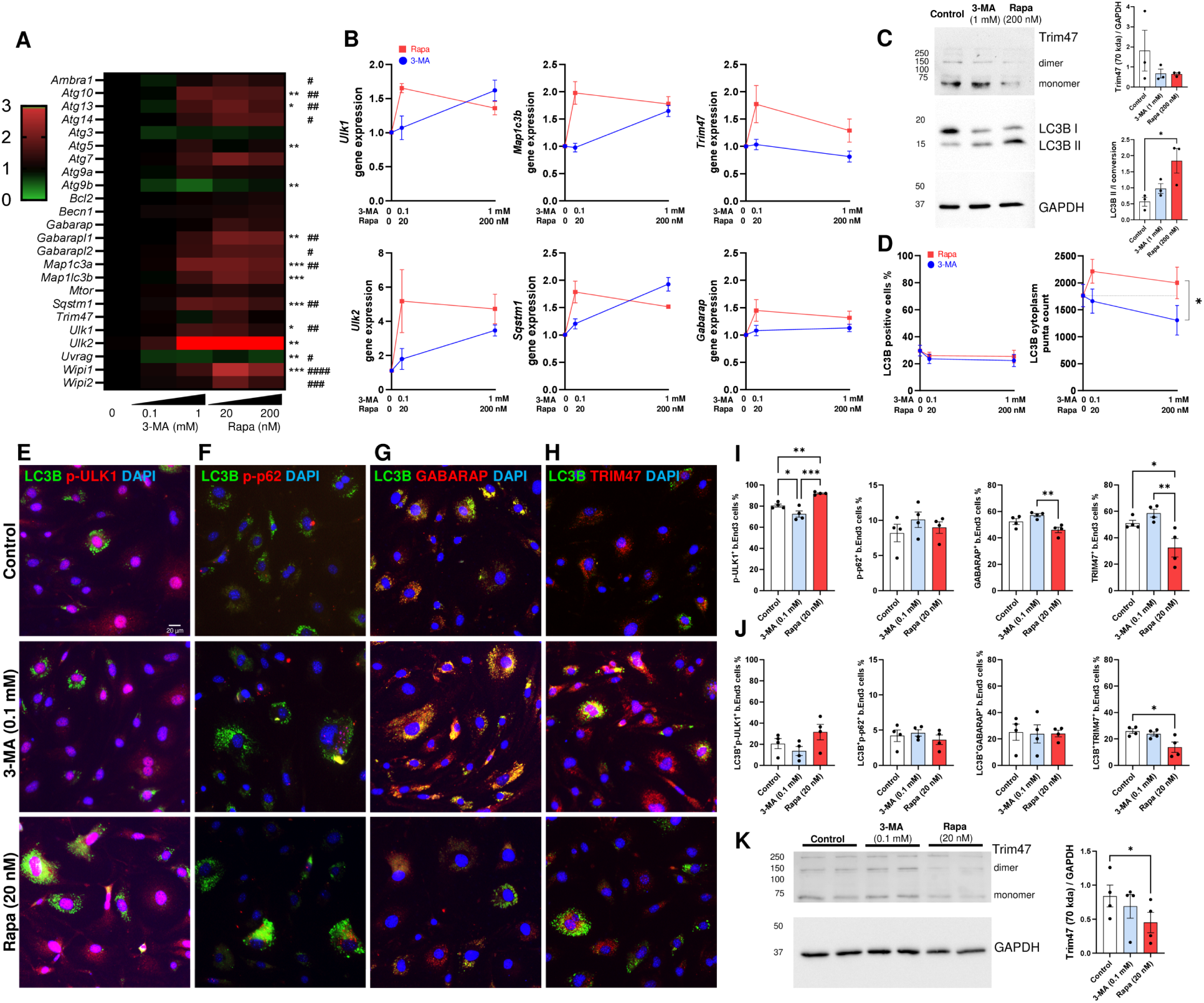
Trim47 expression is modulated by rapamycin-mediated autophagy induction in brain endothelial cells. **A** Heatmap of gene expression across autophagy related genes in bEnd.3 cells upon autophagy inhibition by 3-MA (0.1, 1 mM) and induction by Rapa (20, 200 nM). The statistical significance of gene expression changes mediated by 3-MA and Rapa were denoted by * and #, respectively. **B** The concentration dependent effects of 3-MA and Rapa were exemplified by *Ulk1*, *Ulk2*, *Map1c3b*, *Sqstm1*/*p62*, *Trim47* and *Gabarap* induced by 3-MA and Rapa. Of note, the high concentration of 3-MA mediated autophagy in the full medium as reported earlier (Wu et al., 2010). **C** Immunoblot showing Trim47 expression at both the monomer (70 kDa) and dimer (∼140 kDa) form were reduced by autophagy induction at high concentration of 3-MA (1 mM) and Rapa (200 nM) indicated by the significant LC3B conversion. **D-G** Representative immunocytochemistry image showing the colocalization between LC3B with p-ULK1^Ser555^, p-p62^Ser349^ GABARAP and Trim47, respectively. **H** Quantification of the LC3B^+^ cells and LC3B^+^ puncta in bEnd.3 cells showed a significant difference between 3-MA and Rapa treatment. **I** Quantification of p-ULK1^Ser555^, p-p62^Ser349^ GABARAP and Trim47-expressing cells upon 3-MA and Rapa treatment. **J** Quantification of colocalization between LC3B with p-ULK1^Ser555^, p-p62^Ser349^ GABARAP or Trim47 showed a significant reduction of Trim47-LC3B double positive cells upon Rapa treatment. **K** Immunoblot showing the reduction of Trim47 monomer (70 kDa) by low concentration of Rapa (20 nM), but not 3-MA (0.1 mM), as in the immunocytochemistry. One-way ANOVA repeated measures followed by Tukey’s multiple comparisons test (* or #, p < 0.05; ** or ## P<0.01, *** or ###, P<0.001, ####, P<0.0001).

The bioinformatic and *in silico* analysis thus far suggested that TRIM47 is selectively expressed by ECs in the brain and is a plausible partner of LC3B during autophagic conditions. We experimentally tested these findings on a murine brain-derived endothelial cell line, bEnd.3, using a pharmacological approach. By applying 3-methyladenine (3-MA; 0, 0.1, 1 mM) and rapamycin (Rapa, 0, 20, 200 nM) in full medium for 24 h, we modulated autophagy inhibition and activation on bEnd.3, respectively (Cheng et al., 2020; Wu et al., 2010). At transcription level, 3-MA and Rapa induced differential expression of autophagy genes at the low concentration, such as *Ulk1*, *Ulk2*, *Map1c3b* (LC3B), *Sqstm1* (p62) and *Gabarap* (Fig. 4A, B). At low concentration, Rapa increased *Trim47* transcription but did not reach statistical significance while 3-MA had no effects. Of note, the pleiotropic effects of 3-MA at different concentration were observed as reported (Wu et al., 2010). Immunoblotting showed that Rapa (200 nM) induced a significant increase of LC3B II/LC3B I conversion indicative of autophagy, and such induction was associated with a decrease of Trim47 (Fig. 4C).

To visualize the Trim47 expression in association with different stages of autophagy, the treated cells were examined for LC3B colocalization with p-ULK1^Ser555^, p-p62^Ser349^, GABARAP and Trim47, respectively (Fig. 4D-H). LC3B is the ATG8 member that serves as the marker of autophagy process. The phosphorylation of ULK at serine 555 is mediated by AMPK and is a critical for starvation-induced autophagy at the initiation stage (Egan et al., 2015). The phosphorylation of p62/SQSTM1 at serine 349 is induced selective autophagy (Ichimura et al., 2013). GABARAP is an ATG8 member that is known to associate with LC3B that regulates ULK1 activity in autophagosome formation in response to starvation (Grunwald et al., 2020). The inhibition and promotion of autophagy by 3-MA and Rapa did not alter the overall percentage of LC3B^+^ bEnd.3 cells, but quantifications showed that the two compounds mediated opposing effects on cytoplasmic LC3B puncta count per field (P < 0.05, Fig. 4D). Further quantifications of p-ULK1^Ser555+^ cells showed that the percentage of bEnd.3 at initiation stage was significantly reduced by 3-MA (by 10.1%, 0.1 mM, P < 0.05) but increased by Rapa (by 14%, 20 nM, P < 0.01, Fig. 4I). A 2.3-fold difference in LC3B-p-ULK1^Ser555^ double positive cells was found between 3-MA and Rapa treatment, suggesting differential levels of autophagy induction, as expected (Fig. 4J). In these conditions, the number of p-p62^Ser349^ positive and LC3B-p-p62^Ser349^ double positive bEnd.3 cells remained unchanged, suggestive of minimal selective autophagy. The bEnd.3 cells expressing GABARAP, an ATG8 family member that mediates opposing function against LC3 (Grunwald et al., 2020), was significantly lowered by Rapa compared with 3-MA (20%, P < 0.01). Finally, immunocytochemistry showed that Trim47 expression appeared as puncta and was localized both in the cytoplasm and nucleus (Fig. 4H and Fig. 5). While autophagy inhibition by 3-MA increased Trim47^+^ bEnd.3 cells by 15.3% (vs control, autophagy activation by Rapa reduced Trim47^+^ bEnd.3 cells by 36.1% (vs control, P < 0.05) (Fig. 4I). Of note, the significant difference in Trim47^+^ bEnd.3 cell numbers between 3-MA and Rapa treatments appeared as an opposing trend to p-ULK1^Ser555^ induction. The reduction of Trim47 monomer (70 kDa) by low concentration of Rapa (20 nM), but not 3-MA (0.1 mM), found in the immunocytochemistry was confirmed by immunoblotting (Fig. 4K). As the colocalization between Trim47 and LC3B (Fig. 4D) was highly comparable to that between GABARAP and LC3B (Fig. 4F), and such Trim47^+^ LC3B^+^ double positive cells were significantly reduced after Rapa treatment (Fig. 4G), these data suggested that Trim47 is closely associated with LC3B as predicted by the *in silico* model and it may be a negative regulator of autophagy.

**Figure 5.**
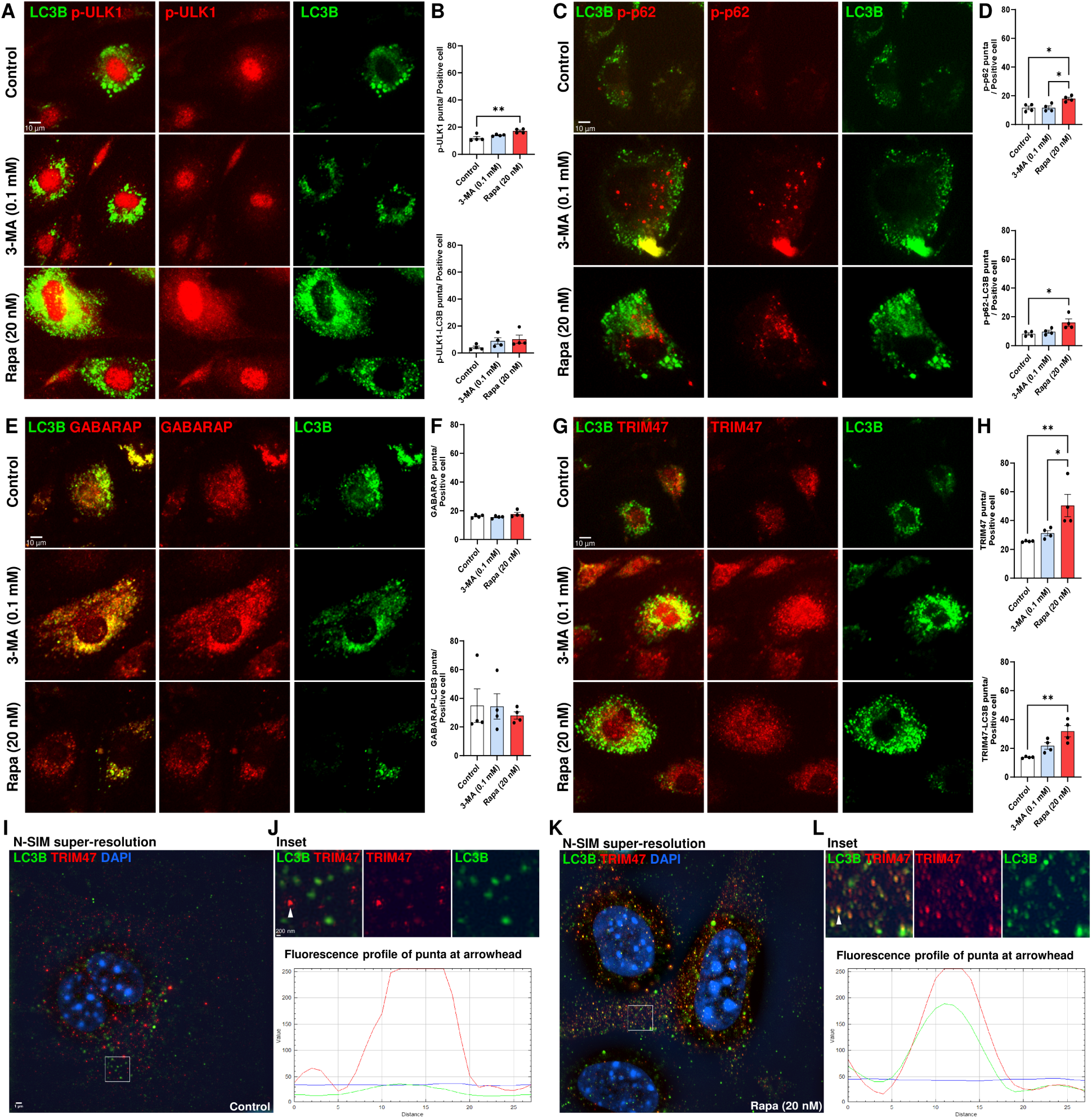
Trim47 colocalization with LC3B in autophagic brain endothelial cells. Representative immunocytochemistry image and quantification of **A, B** LC3B-p-ULK1^Ser555^, **C, D** LC3B-p-p62 (p-p62^Ser349^), **E, F** LC3B-GABARAP, and **G, H** LC3B-Trim47 colocalization in control, 3-MA, or Rapa treatment. **I** Structured Illumination Microscopy (SIM)-based super-resolution image of the LC3B and Trim47 at close proximity under control condition, which the inset area shown at a higher magnification in **J**. The colour profile plot below demonstrates the lack of Trim47-LC3 colocalization in control. **K** SIM-based super-resolution image of the LC3B and Trim47 at close proximity after Rapa treatment (20 nM, 24 h), in which the inset area is magnified in **L**. The colour profile plot below confirmed the colocalization of Trim47-LC3 puncta during autophagy. One-way ANOVA repeated measures followed by Tukey’s multiple comparisons test (*p < 0.05; ** P<0.01). **I, J** and **K, L** share the same scale with scale bar = 1 µm and 200 nm, respectively.

Many TRIM members were experimentally verified to interact with key autophagic adaptors including LC3 (i.e *TRIM5, -17, -20, -21, -22, -28, -41, -49, -55*) or SQSTM1/p62 (i.e *TRIM5, -13, -17, -21, -22, -28, -49, -50, -55, -63*) and they form TRIM-containing complex during autophagosome formation (Mandell et al., 2014), which serve as platform for macroautophagy and selective autophagy (Kimura et al., 2016). To examine such potential interactions, we examined the puncta colocalization between LC3B with known adaptors including p-ULK1 (p-ULK1^Ser555^) p62 (p-p62^Ser349^), GABRARAP and Trim47 in bEnd.3 cells with or without 3-MA or Rapa treatment (Fig. 5). The increased p-ULK1^+^ cells were associated with an increase of p-ULK1^+^ cytoplasmic puncta (Fig. 5A, B). Although the number of p-p62^+^ cell remain unchanged, the p-p62 and p-p62-LC3B^+^ double positive puncta per cell was significantly increased by Rapa (P < 0.05, Fig. 5C, D). For GABARAP, a known ATG8 member and an LC3B partner, the average number of GABARAP^+^ and GABARAP^+^LC3B^+^ puncta in each positive cell remained unchanged by 3-MA and Rapa treatment (Fig. 5E, F). Finally, a clear colocalization between Trim47^+^ and LC3B puncta was identified (Fig. 5G, H). Importantly, although the cell number of Trim47^+^ LC3B^+^ b.End3 was significantly reduced by Rapa (Fig. 4J), the number of Trim47^+^ and Trim47^+^LC3B^+^ puncta in the persisting Trim47-expressing bEnd.3 cells were doubled by Rapa treatment (Fig. 5F). Using SIM-based super-resolution microscopy (Fig. 5I-L), we confirmed the close proximity between LC3B and Trim47 in control conditions (Fig. 5I) with their colocalization at a resolution < 200 nm in Rapa-mediated autophagy (Fig. 5K).

The downregulation of *Trim47* autophagic conditions of ECs and the reduction of Trim47-expressing cells upon Rapa treatment suggested that Trim47 may be negatively correlated with autophagy. We proposed that TRIM47 maybe one of the negative regulators of autophagy among the TRIM family members (Mandell et al., 2014). To test, we silenced Trim47 (siTrim47) in bEnd.3 cells in the presence or absence of 2DG-mediated starvation. In gene expression study, siTrim47 induced a significant differential expression of *Atg7*, *Bcl2 Gabarap*, *Gabarapl1* and *Gabarapl2* (Fig. 6A). While 2DG significantly reduced expression of *Atg7* and *Gabarap* family, silencing Trim47 in bEnd.3 cells doubled this expression in all conditions (Fig. 6B). *Atg7* is an essential regulator of cytoplasmic to vacuole transport in autophagy while the *Gabarap* family is important for the formation of late autophagosome, and these suggested that the loss of Trim47 may disinhibit autophagy, especially at the later stage. Indeed, in immunocytochemistry, gene silencing of *Trim47* augmented the expression of p-p62^Ser349^, p-ULK1^Ser555^ and GABARAP in bEnd.3 cells independent of 2DG-mediated starvation (Fig. C-E). Neither siTrim47 nor 2DG altered the number of p-ULK^+^ cells but siTrim47 significantly increased the number of p-ULK1^Ser555+^ in each positive cell (P = 0.0097, Fig. 6F). While only 2DG significantly induced p-p62 expressing cells (P = 0.0028) and p-p62 puncta per cell (P = 0.0139), siTrim47 induced the highest number of p-p62 puncta per cell (vs Control, P = 0.0424, Fig. 6G). Silencing Trim47 also significantly increased the percentage of GABARAP-expressing cells (P = 0.0009), but the number of GABARAP puncta was unaffected by either treatment (Fig. 6H). The examination of LC3B showed that siTrim47 significantly increased the number of LC3B^+^ bEnd.3 cells with or without 2DG treatment (Fig. 6I). Importantly, the loss of Trim47 also increased the percentage of cells with vacuole formation (Fig. 6J, K). Quantifications showed that siTrim47 increased LC3B^+^ b.End3 cells a 2.2-fold and 2.8-fold in control and 2DG treatment, respectively (Fig. 6L). Although siTrim47 did not increase the number of LC3B^+^ puncta, it significantly increased the number of vacuole positive cells by 4.8 and 7.6 folds in control and 2DG treatment, respectively (Fig. 6M). Together, these data indicate that the loss of Trim47 may disinhibit autophagy in brain ECs and suggested that Trim47 is an endogenous inhibitor of autophagy.

**Figure 6.**
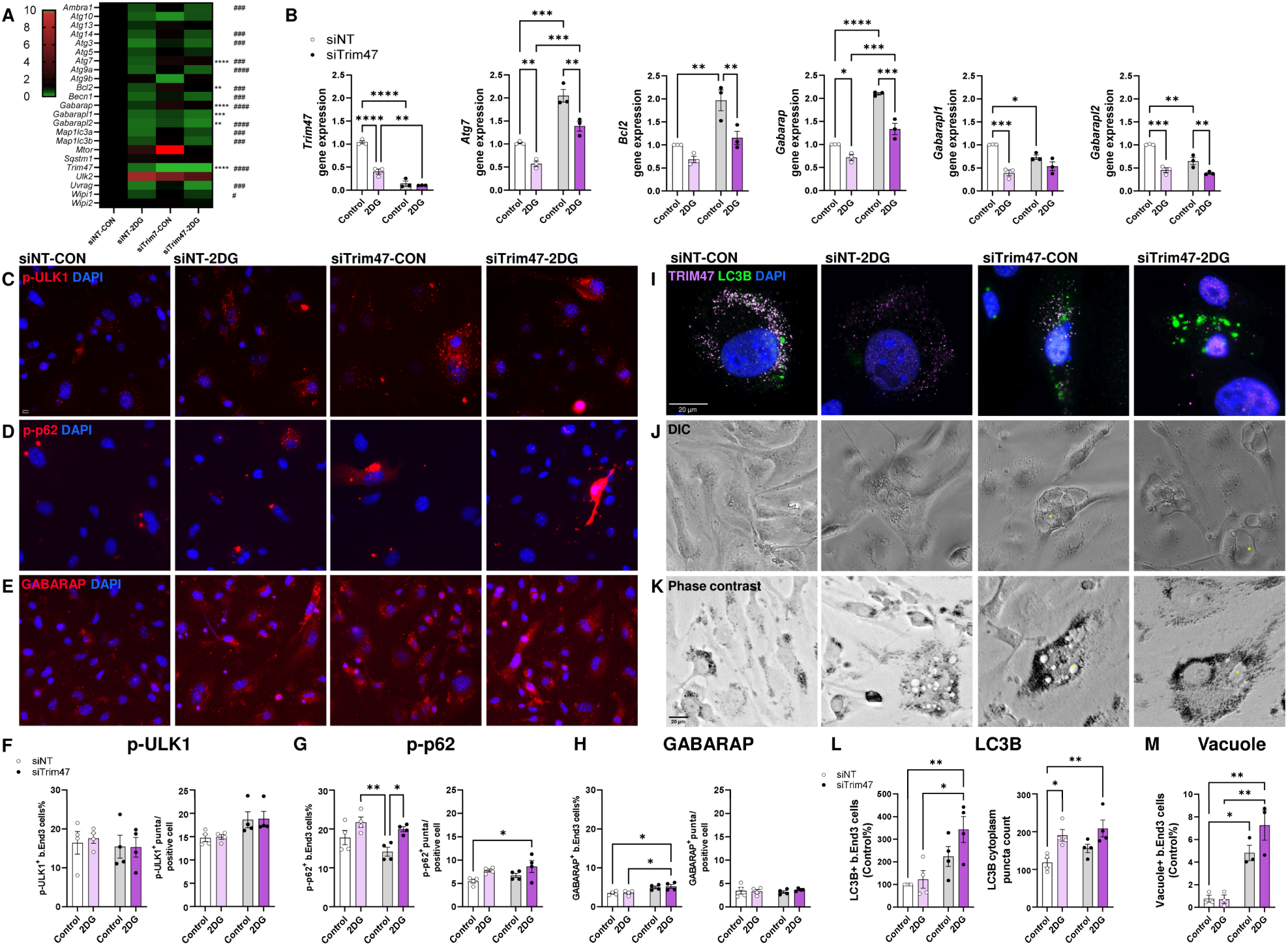
Silencing Trim47 increased autophagy in brain endothelial cells. **A** Heatamp of gene expression analysis of b.End3 cells with non-targeted gene silencing (siNT) and siTrim47 with or without 2DG-mediated starvation Two-way ANOVA repeated measures followed by Holm-Šídák’s multiple comparisons test (*p < 0.05, ** P<0.01, ***P<0.001, ****P<0.0001). **B** Individual gene expression analysis of *Trim47, Atg7, Bcl2, Gabarap, Gabarapl1* and *Gabarapl2* demonstrating that the loss of *Trim47* disinhibit autophagy. **C-E** Representative immunocytochemistry image of bEnd.3 cells expressing p-ULK1^Ser555^, p-p62^Ser349^ and GABARAP treated by siNT or siTrim47 with or without 2DG treatment. **F-H** Quantifications of p-ULK1^Ser555^, p-p62^Ser349^ and GABARAP expressing upon siTrim47 with or without 2DG starvation were shown, where **F** silencing Trim47 significantly increased the number of p-ULK1^Ser555+^ puncta in each positive cell (P = 0.0097), **G** induced the highest number of p-p62 puncta per cell (vs Control, P = 0.0424) and **H** significantly increased the percentage of GABARAP-expressing cells (P = 0.0009). **I** Representative immunocytochemistry image showing the effect of siTrim47 and 2DG on LC3B and Trim47 expression in b.End3 cells. **J, K** Differential interference contrast (DIC) and phase contrast of bEnd.3 cells showing vacuole formation (*) under siTrim47 conditions, respectively. **L, M** Quantification of LC3B^+^ cells, LC3B^+^ puncta per cell, and vacuole formation showing that sip-pTrim47 significantly increases autophagy responses in bEnd.3 cells. Two-way ANOVA followed by Tukey multiple comparisons test (*p < 0.05, ** P<0.01).

Among the SNPs of *TRIM47* at 17q25 that leads WMH, only one missense variant (rs4600514; NP_258411.2; p.R187W) has potentially damaging consequences (Mishra et al., 2022). It remains unknown whether this mutation will lead to the loss-of-function or gain-of toxicity-effects on brain ECs. To investigate, we tracked the location of R187 in the predicted 3D structure of human TRIM47 and mouse Trim47 protein (equivalent to R191) (Fig. 7A, B). In both models, the location of this residue is situated in the RING-finger domain which is responsible for E3 ubiquitin ligase function. However, this residue is far from the LIR motif (NQWEQL or SQWEQL) that is responsible for autophagy function. Yet, TRIM family members are known to form homodimer, heterodimer and oligomer which may in turn contribute to the recognition of diverse substrates and catalytic regulations (Koliopoulos et al., 2016; Koliopoulos et al., 2018). We asked if the TRIM47 model may form the anti-parallel coiled-coil dimerization as in TRIM25 (Koliopoulos et al., 2018; Sanchez et al., 2014), and whether the residue at R187 may have any implications on the dimerization (Fig. 7C-E). The *in silico* simulation showed that it is likely that TRIM47 may form coiled-coil dimerization as in TRIM25, but the RING-finger domain remains distant from the dimerization sites (Fig. 7C - E). To validate the TRIM47 dimerization, we resolved proteins isolated from bEnd.3 cells treated with 2DG in PAGE without sodium dodecyl sulfate denaturation. A significant increase of potential TRIM47 dimers at approximately 140 kDa (143.4 ± 2.5 kDa) and 130 kDa (134.6 ± 2.3 kDa) were identified. In such non-denaturing condition, a putative oligomer above 250 kDa was also observed (Fig. 7F). Given the size of Trim47 monomer is 70 kDa, these findings suggested that Trim47 may form homodimer or heterodimer. In addition, the potential formation of coiled-coil dimerization is the basis of the formation of a hexagon mesh network in TRIM5/TRIM5α which bridges the RING domains between two molecules and provide a scaffold as an oligomer for autophagy (Carter et al. 2020; Mandell et al. 2014). The dimerization and putative oligomerization of TRIM47 suggested that a similar oligomerization is plausible (Fig. 7G).

**Figure 7.**
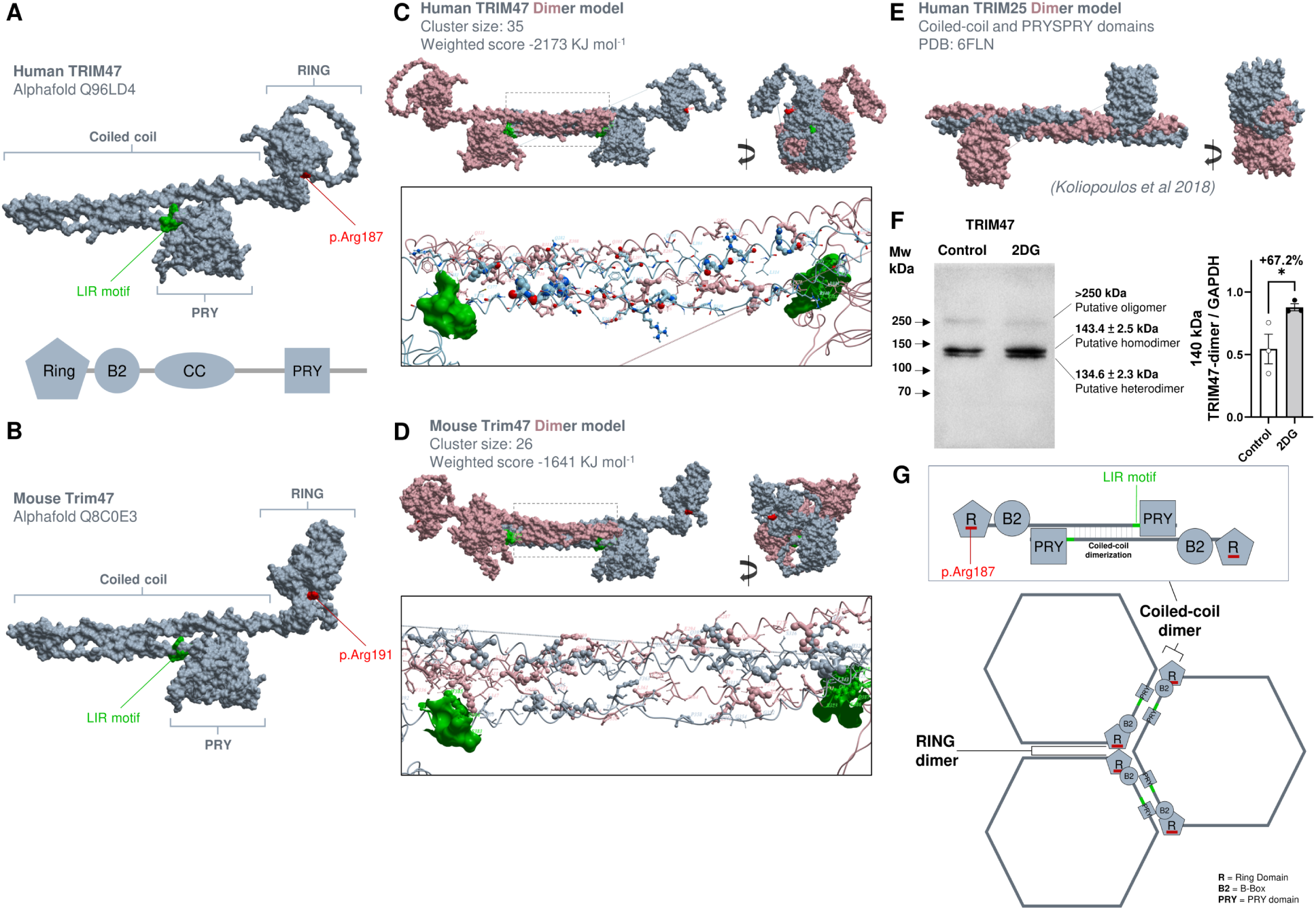
WMH associated SNP is located at the RING region of TRIM47 at distance from LIR motif. The *in silico* model demonstrating **A** human TRIM47 protein and **B** mouse Trim47 structure with the location of the LIR motif in green and the missense variant p.Arg187 (p.Arg191 in mouse) shown in red. **C, D** ClusPro-based simulation of coiled-coil homodimer of human and mouse TRIM47 model demonstrating the distance between two LIR motif (green) and the lack of interactions between the missense variant p.Arg187 (red) in the RING domain. **E** Reproduced TRIM25 coiled-coil homodimer model described by Koliopoulos et al 2018. F Immunoblotting of bEnd.3 lysate in control and 2DG-treatment resolved under non-denaturing conditions showing the increase of Trim47 dimers at ∼140 kDa (143.4 ± 2.5 kDa) and 130 kDa (134.6 ± 2.3 kDa), respectively. The 140 kDa dimer is likely a homodimer of Trim47 (monomer = 70 kDa), while the 130 kDa dimer could be a heterodimer with other Trim family members. An additional band at above 250 kDa suggested that the Trim47 might form oligomers. **G** The formation of TRIM47 coiled-coil homodimers could be the basic building block that may allow the formation of an oligomer as a hexagon mesh of network that may be part of the scaffold (TRIMosome) to facilitate autophagy as earlier described in TRIM5/TRIM5α (Carter et al. 2020; Mandell et al. 2014).

## Discussion

WMH formation is highly correlated with cognitive decline in aging and the progression of age-related dementia. The common identification of genetic risk locus at 17q25 found in GWAS studies has provided clues towards the molecular mechanism of WMH formation (Fornage et al., 2011; Persyn et al., 2020; Traylor et al., 2019; Verhaaren et al., 2015). Here, we proposed that *TRIM47*, one of the two TRIM family members at this genetic locus, is selectively expressed in the brain ECs and plays a role in autophagy. We identified the potential binding sites with the key autophagy adaptor LC3B *in silico* and found that TRIM47 is a negative regulator of autophagy *in vitro*. Autophagy is the key process for ECs remodeling during hypertension flux (Vion et al., 2017), and, in turn, hypertension is a major physiological risk factor for WMH formation (Chesebro et al., 2020; Hajjar et al., 2011; Verhaaren et al., 2013). Our results support that dysfunctional TRIM47 disrupts brain ECs autophagy which might lead to WMH formation.

The 80 plus members of the TRIM family not only serve as E3 ubiquitin ligases, but also plays a role in autophagy. In a systemic knockdown screening of 67 human *TRIM* in HeLa cells, it was found that TRIM members can be positive (TRIM3, -5, -6, -22, -23, -28, -31, -34, -38, -41, -42, -44, -45, -48 to -50, -55, -58, -59, -61, -73, and -74) or negative (TRIM1, -2, -10, -17, -26, -33, -51, -65, -72, and -76) regulators of autophagy at different stages (Di Rienzo et al., 2020; Mandell et al., 2014). Our hypothesis that TRIM47 dysfunction may lead to EC pathology during WMH formation is supported by a recent study showing that the knockdown of TRIM47 resulted in an increase of EC permeability *in vitro* (Mishra et al., 2022). Nevertheless, this study did not address the cellular mechanism connecting the TRIM47 loss-of-function and permeability change. The two key cellular homeostasis processes important for EC functions, ubiquitination and autophagy (Majolee et al., 2019; Meng et al., 2022), are both regulated by TRIMs. TRIM47 was showed to regulate TNFα-mediated lung EC activation by degrading TRAF2 through K63 ubiquitination (Qian et al., 2022), but its role in autophagy is unknown. Here, we confirmed that TRIM47 is selectively expressed in the ECs of human and mouse brain, and we demonstrated that *TRIM47* is downregulated among the relevant models of hypertension in EC with ongoing autophagy. Of note, we observed that *Trim47* transcription shifts occur acutely (days) upon experimental ischemic models (see Table S3) and returns to normal in chronic models (weeks). Indeed, in our recent report, we detected minimal *Trim47* transcription change in the mouse corpus callosum after a four-weeks bilateral carotid artery stenosis model simulating subcortical ischemic vascular dementia (Takase et al., 2021). These observations suggest that TRIM47 may regulate autophagy at the early stage of cerebral ischemia. TRIM family members regulate autophagy at different stages by targeting adaptor proteins such as AMPK, ULK1 or Beclin1, it is also proposed that these proteins form a TRIM-containing complex during autophagosome formation (Di Rienzo et al., 2020). As in TRIM5α, a positive regulator of autophagy (Mandell et al., 2014), we identified the LIR motif (NEWEQL) in TRIM47 and its potential binding partner residues in LC3B using *in silico* and such interaction was visualized by the super resolution microscopy.

The putative interaction with LC3B may confer the autophagy regulatory function to TRIM47. In bEnd.3 cells, autophagy inhibition mediated by 3-MA and autophagy activation mediated by rapamycin were associated with Trim47 upregulation and downregulation, respectively. This opposing expression pattern suggested that Trim47 may regulate autophagy in EC. The gene silencing experiment confirmed that Trim47 is a negative regulator of autophagy in brain ECs. Other TRIM family members negatively regulate autophagy by targeting different key autophagic factors such as TFEB (Wang et al., 2018), AMPK (Pineda et al., 2015) or Beclin1 (Han et al., 2018) through proteasome degradation, transcription or microRNA regulations (Di Rienzo et al., 2020). These regulatory functions of TRIM protein are highly associated with their intracellular distribution, for instance, TRIM52 regulate immune functions by acting in the nucleus (Fan et al., 2017) while TRIM20 regulates E3 ligase functions in the cytoplasm (Hatakeyama, 2017). The dual localization of Trim47 suggested that it may serve autophagy regulatory functions in the nucleus and cytoplasm of brain ECs. Silencing Trim47 not only resulted in a significant upregulation of players (*Atg7 Bcl2* and *Gabarap*) important for in autophagosome formation (Collier et al., 2021; Ma et al., 2013), but also increased the ECs with phosphorylation of ULK1 and p62 as well as vacuole formation. As ULK-mediates the phosphorylation of p62 (Ikeda et al., 2023), Trim47 is therefore pivotal to both early and later stage of autophagy (Egan et al., 2015). TRIM65, the other TRIM member at locus 17q25, is known to inhibit autophagy also by controlling *Atg7* expression in lung cancer cells (Pan et al., 2019). It remains obscure whether the TRIM47 and TRIM65 mediate autophagy regulation by jointly targeting *Atg7* in brain ECs, but it warrants further investigation.

The possible TRIM47 the pathogenic variant (rs600514, missense, NP_258411.2: p.Arg187Trp) is located in the RING domain and may have a direct implication on the E3 ligase activity. Yet, TRIM members are known to form homodimer, heterodimer and oligomer in order to function (Hatakeyama, 2017) (see Fig. 7G), which opens up other possible mechanisms by which this particular variant might increase the risk of WMH. First, the self-association of two TRIM monomers is enabled by the highly conserved coiled coil which forms an interdigitating anti-parallel helical hairpin (Sanchez et al., 2014). Such a homodimer is proposed to be the building block of more complex oligomers with the TRIM5/TRIM5α being the best studied example. Second, the dimerization between the RING domain is critical for E3 ligase activity and is conserved among TRIM subfamily C-IV where TRIM47 belongs (Fiorentini et al., 2020). Finally, as exemplified by TRIM5/TRIM5α, these antiparallel, coiled-coil homodimers with the RING domain at both ends may form a hexagon scaffold (TRIMosome) to facilitate autophagy (Carter et al., 2020; Mandell et al., 2014). Our in silico modeling shows that TRIM47 can form such a dimer and our in vitro work shows that a dimeric form of TRIM47 is increased when autophagy is induced. Although the distance (194 amino acids) between the RING domains and LIR motif at the PRY domain would seem to make the missense variant more relevant to ubiquitination than to autophagy, the dimerization or oligomerization of TRIM47 may impact autophagy as well as E3 ligase activity. TRIM family members are most likely to form heterodimers with structurally related members (Sanchez et al., 2014). We note that TRIM47 and TRIM65, both encoded by genes at 17q25 locus are both members of the same subfamily, C-IV (Hatakeyama, 2017). This leads us to speculate that the two proteins may be coordinately regulated during normal physiological functions and form heterodimers specifically targeted to autophagic functions. The rs600514 variant may create a pathogenic heterodimer leading to improper autophagy and thus to WMH formation.

Brain vasculature is a highly dynamic system, where flexibility is conferred by the process of autophagy in the ECs that form and important part of its structure. Our findings support a model in which dysfunctional TRIM47 contributes to autophagy failure in brain ECs and in this way compromises the remodeling of the cerebrovasculature, especially in conditions favoring hypertension. The impaired responsiveness to the normal (or enhanced) shear stresses that results confers an elevated risk of WMH development and offers a satisfying mechanistic explanation for the link between the rs600514 TRIM47 missense mutation and the increased risk of WMH.

## DECLARATIONS

### Ethics approval and consent to participate

- This study involves human tissue

`o` All human tissue experiments from the brain bank in the Neuropathology Core of Alzheimer’s Disease Research Center (ADRC) at University of Pittsburgh Medical Center with approvals from the Committee for Oversight of Research and Clinical Training Involving Decedents at University of Pittsburgh. The ADRC is funded by NIH P30 AG066468-02. This cohort of postmortem tissue from the brain bank is ruled as exempt by IRB at University of Pittsburgh Research as it only involves the collection or study of existing data, documents, records, pathological specimens or diagnostic specimens, if these sources are publicly available or if the information is recorded by the investigator and in such a manner that subjects cannot be identified, directly or through identifiers linked to the subjects.
- This study involves animals

`o` All animal experiments in this study are approved by Animal Subjects Ethics Sub-committee (ASESC), Research Committee, The Hong Kong Polytechnic University with ASESC reference number 20-21/167-HTI-R-GRF

### Consent for publication

- Not applicable

### Availability of data and material

- The datasets during and/or analyzed during the current study available from the corresponding author on reasonable request.

## Acknowledgments

- The present work is generously supported by General Research Fund from Research Grant Council (GRF15101422), Hong Kong Special Administrative Region. Dr. Kai-Hei Tse is also supported by PolyU Start-up Fund P0030307. The initial concept of the present work was funded by The Leo and Anne Albert Charitable Trust to Prof. Karl Herrup, Prof. Myriam Fornage, Dr Ken Arai and Dr Kai-Hei Tse
- Prof. Karl Herrup is supported by Start-up Fund at Department of Neurobiology, University of Pittsburgh and the Australian National Health and Medical Research Fund (APP1160691), the Pennsylvania Department of State (4100087331) and additional support from the NIA (R01 AG069912).
- Dr Kofler and Alzheimer’s Disease Research Center at the University of Pittsburgh are supported by NIA P30 AG066468 and NIA P50 AG005133.

## Authors’ contributions

- S.H.S.Y performed cell culture experiments, immunohistochemistry and imaging. R.H.S.L performed image analysis. K.H.T performed RNA-seq meta-analysis and in silico modeling simulations. G.W.Y.C and I.W.T.M. performed animal husbandry. P.H performed the histopathology test. C.K., F.M., K.H., J.K, performed immunohistochemistry on human tissue. S.H.S.Y and K.H.T wrote the first draft of the manuscript. K.H., M.F., K.A., and K.H.T conceptualized the study, designed the experiment, and edited the manuscript. All authors read, critically reviewed, and approved the manuscript. K.H.T supervised the study and obtained funding.

