## Supplementary material for "White matter hyperintensity genetic risk factor *TRIM47* regulates autophagy in brain endothelial cells": Table S1

| **Target** | **Forward** | **Reverse** |
|---|---|---|
| *Ambra1* | CCAGAGAAGAATGCTGTACGAAT | TCCATCGAGTCTTATCCTCCAC |
| *Atg10* | GTAGTTACCAAGTGCCGGTTC | AGCTAACGGTCTCCCATCTAAA |
| *Atg12* | TCCCCGGAACGAGGAACTC | TTCGCTCCACAGCCCATTTC |
| *Atg13* | CCAGGCTCGACTTGGAGAAAA | AGATTTCCACACACATAGATCGC |
| *Atg14* | GAGGGCCTTTACGTGGCTG | AATAGACGAAATCACCGCTCTG |
| *Atg3* | ACACGGTGAAGGGAAAGGC | TGGTGGACTAAGTGATCTCCAG |
| *Atg5* | TGTGCTTCGAGATGTGTGGTT | GTCAAATAGCTGACTCTTGGCAA |
| *Atg7* | GTTCGCCCCCTTTAATAGTGC | TGAACTCCAACGTCAAGCGG |
| *Atg9a* | CAGTTTGACACTGAATACCAGCG | AATGTGGTGCCAAGGTGATTT |
| *Atg9b* | ATGTACCCGAAGGACTCCG | CATTCCGCTGATGATAGCTGT |
| *Bcl2* | ATGCCTTTGTGGAACTATATGGC | GGTATGCACCCAGAGTGATGC |
| *Becn1* | ATGGAGGGGTCTAAGGCGTC | TCCTCTCCTGAGTTAGCCTCT |
| *Gabarap* | AAGAGGAGCATCCGTTCGAGA | GCTTTGGGGGCTTTTTCCAC |
| *Gabarapl1* | GGACCACCCCTTCGAGTATC | CCTCTTATCCAGATCAGGGACC |
| *Gabarapl2* | TCGGGCTCTCAGATTGTTGAC | ATGGCCTTCTCGGAGGGAA |
| *Map1lc3a* | GACCGCTGTAAGGAGGTGC | CTTGACCAACTCGCTCATGTTA |
| *Map1lc3b* | TTATAGAGCGATACAAGGGGGAG | CGCCGTCTGATTATCTTGATGAG |
| *Mtor* | ACCGGCACACATTTGAAGAAG | CTCGTTGAGGATCAGCAAGG |
| *Sqstm1* | AGGATGGGGACTTGGTTGC | TCACAGATCACATTGGGGTGC |
| *Trim47* | GAGATCATCGAAGGATGGGTCA | CCCGCAGAACCAGACTGAG |
| *Ulk1* | AAGTTCGAGTTCTCTCGCAAG | CGATGTTTTCGTGCTTTAGTTCC |
| *Ulk2* | AGCTTCAGCATGAAAACATCGT | CGATTGGCATAAGACAACAGGA |
| *UVRAG* | ACATCGCTGCTCGGAACATT | CTCCACGTCGGATTCAAGGAA |
| *Wipi1* | CTGCTTCTCTTTCAACCAAGACT | ACGTCAGGGATTTCATTGCTT |
| *Wipi2* | AGGATAACACGTCCCTAGCTG | TCTCTCCACAATGCAGACATCT |

Table S1 – Primer Sequence for qPCR
