## Supplementary material for "White matter hyperintensity genetic risk factor *TRIM47* regulates autophagy in brain endothelial cells": Table S2

| Analysis | Platform | GSE Number | Sample ID | Sample for comparison | Gene Symbol | Log Fold Change | log CPM | P Value | FDR /adjusted P Value | Power | BCOV |
|---|---|---|---|---|---|---|---|---|---|---|---|
| Human - Aorta EC Shear Stress | GEO2R | GSE199709 | GSM5982477<br>GSM5982478<br>GSM5982479<br>GSM5982480<br>GSM5982481<br>GSM5982482 | HAEC_0dy_0h_1<br>HAEC_0dy_0h_2<br>HAEC_0dy_0h_3<br>HAEC_12dy_12h_1<br>HAEC_12dy_12h_2<br>HAEC_12dy_12h_3 | *TRIM47* | 0.410306 | | 0.00144 | 0.00964 | | |
| Human - Aorta EC Shear Stress | GEO2R | GSE199709 | GSM5982477<br>GSM5982478<br>GSM5982479<br>GSM5982483<br>GSM5982484<br>GSM5982485 | HAEC_0dy_0h_1<br>HAEC_0dy_0h_2<br>HAEC_0dy_0h_3<br>HAEC_after_24h_1<br>HAEC_after_24h_2<br>HAEC_after_24h_3 | *TRIM47* | 0.164828 | | 1.25E-01 | 3.08E-01 | | |
| Human - EC Autophagy TFEB | GEO2R | GSE108384 | GSM2897074<br>GSM2897075<br>GSM2897076<br>GSM2897077 | Ad-GFP treated-rep1<br>Ad-GFP treated-rep2<br>Ad-TFEB treated-rep1<br>Ad-TFEB treated-rep2 | *TRIM47* | -1.1642143 | | 0.00000514 | 0.000328 | | |
| Mouse - MCAO cortex | GREIN | GSE131712 | GSM3814504<br>GSM3814505<br>GSM3814506<br>GSM3814510<br>GSM3814511<br>GSM3814512 | Sham_C1<br>Sham_C2<br>Sham_C3<br>R24h_C1<br>R24h_C2<br>R24h_C3 | *Trim47* | -3.163 | 2.69 | 4.9475E-16 | 4.51735E-14 | 1 | 0.279 |
| Mouse - MCAO cortex | GREIN | GSE131712 | GSM3814504<br>GSM3814505<br>GSM3814506<br>GSM3814516<br>GSM3814517<br>GSM3814518 | Sham_C1<br>Sham_C2<br>Sham_C3<br>R28d_C1<br>R28d_C2<br>R28d_C3 | *Trim47* | 0.467 | 0.596 | 0.346148286 | 0.902496748 | 0.092 | 0.357 |
| Mouse - MCAO endothelial cell | GREIN | GSE122345 | GSM3464400<br>GSM3464406<br>GSM3464408<br>GSM3464414<br>GSM3464399<br>GSM3464405<br>GSM3464407<br>GSM3464413 | A2_control contralateral<br>D2_control contralateral<br>E2_control contralateral<br>H2_control contralateral<br>A1_control ipsilateral<br>D1_control ipsilateral<br>E1_control ipsilateral<br>H1_control ipsilateral | *Trim47* | 0.186 | 8.662 | 0.504908879 | 0.845924894 | 0.152 | 0.273 |
| Mouse - Pulmonary Hypertension | GREIN | GSE180169 | GSM5454559<br>GSM5454560<br>GSM5454561<br>GSM5454562<br>GSM5454563<br>GSM5454564<br>GSM5454565<br>GSM5454566<br>GSM5454567 | RNA-seq of TdTomato+ cells from Cont1 mouse<br>RNA-seq of TdTomato+ cells from Cont2 mouse<br>RNA-seq of TdTomato+ cells from Cont3 mouse<br>RNA-seq of TdTomato+ cells from Cont4 mouse<br>RNA-seq of TdTomato+ cells from Cont5 mouse<br>RNA-seq of TdTomato+ cells from PAH1 mouse<br>RNA-seq of TdTomato+ cells from PAH2 mouse<br>RNA-seq of TdTomato+ cells from PAH3 mouse<br>RNA-seq of TdTomato+ cells from PAH4 mouse | *Trim47* | -1.848 | 5.636 | 0.000141208 | 0.009195056 | 1 | 0.47 |

**Table S2. Details of Gene Ontology analysis of RNA-seq dataset from experiments of autophagy or hypertensive conditions**

EC, endothelial cells; PAH, pulmonary arterial hypertension; MCAO, Middle Cerebral Artery Occlusion; GO, gene ontology; n.d., non-detectable; FDR, false discovery rate ; BOCV, biological coefficient of variation
