## Supplementary material for "White matter hyperintensity genetic risk factor *TRIM47* regulates autophagy in brain endothelial cells": Table S3

| GEO accession | Publication | Species | Model | Comparison | Time | DEG directions | Gene Ontology (GO) biological process complete | GO ID | GO Ref Gene# | Mapped Gene# | Expected Gene# | Fold Enrichment | Raw P value | FDR |
|---|---|---|---|---|---|---|---|---|---|---|---|---|---|---|
| GSE108384 | Fan et al. 2018 | Human | Cell *In vitro* | Ad-GFP HUVEC vs Ad-TFEB HUVEC | 24 h | UP | regulation of autophagy | GO:0010506 | 352 | 77 | 44.95 | 1.71 | 0.00004 | 0.00427 |
| | | | | | | DOWN | negative regulation of macroautophagy | GO:0010507 | 32 | 12 | 5.4 | 2.22 | 0.02810 | 0.49700 |
| GSE199709 | Meng et al. 2022 | Human | Cell *In vitro* | Control aortic ECs vs Shear Stress-treated aortic ECs | 12 h treatment | UP | macroautophagy | GO:0034262 | 198 | 28 | 12.69 | 2.21 | 0.00033 | 0.24200 |
| | | | | | | DOWN | n.d. | n.d. | n.d. | n.d. | n.d. | n.d. | n.d. | n.d. |
| | | | | Control aortic ECs vs Shear Stress-treated aortic ECs | 12 h treatment with 24 h recovery | UP | n.d. | n.d. | n.d. | n.d. | n.d. | n.d. | n.d. | n.d. |
| | | | | | | DOWN | n.d. | n.d. | n.d. | n.d. | n.d. | n.d. | n.d. | n.d. |
| GSE180169 | Rodor et al. 2022 | Mouse | Cell *In vivo* | TdTomato+ EC in Control Lung vs PAH Lung | 3 weeks treatment | UP | protein lipidation involved in autophagosome assembly | GO:0061739 | 6 | 2 | 0.05 | 37.8 | 0.00206 | 0.14000 |
| | | | | | | DOWN | regulation of autophagy | GO:0010506 | 294 | 11 | 5.63 | 1.95 | 0.03300 | 0.61000 |
| GSE131712 | Wu et al. 2019 | Mouse | Tissue *In vivo* | Contralateral MCAO brain cortex vs Ipsilateral brain cortex | 24 h post-injury | UP | negative regulation of macroautophagy | GO:0010507 | 30 | 6 | 1.91 | 3.14 | 0.01900 | 0.32000 |
| | | | | | | DOWN | regulation of autophagy | GO:0010506 | 294 | 50 | 27.43 | 1.82 | 0.00020 | 0.00284 |
| | | | | Contralateral MCAO brain cortex vs Ipsilateral brain cortex | 28 days post-injury | UP | negative regulation of autophagic cell death | GO:1904093 | 5 | 2 | 0.05 | 36.97 | 0.00231 | 0.0453 |
| | | | | | | DOWN | n.d. | n.d. | n.d. | n.d. | n.d. | n.d. | n.d. | n.d. |
| GSE122345 | Wegner et al. 2020 | Mouse | Cell *In vivo* | Contralateral MCAO brain ECs vs Ipsilateral MCAO brain ECs | 72 h post-injury | UP | n.d. | n.d. | n.d. | n.d. | n.d. | n.d. | n.d. | n.d. |
| | | | | | | DOWN | n.d. | n.d. | n.d. | n.d. | n.d. | n.d. | n.d. | n.d. |

**Table S3. Summary of Gene Ontology analysis of RNA-seq dataset from experiments of autophagy or hypertensive conditions**

EC, endothelial cells; PAH, pulmonary arterial hypertension; MCAO, Middle Cerebral Artery Occlusion; GO, gene ontology; n.d., non-detectable; FDR, false discovery rate
