## Supplementary material for "White matter hyperintensity genetic risk factor *TRIM47* regulates autophagy in brain endothelial cells": Table S4

| A. Human TRIM47 (Phyre2 on PDB: 4CG4) x Human LC3B (PDB: 3VTU) | | | |
|---|---|---|---|
| **LIR motif position**: 379-384 **LIR sequence**: NQWEQL | | | |
| | **Balanced** | **Electrostatic** | **Hydrophobic** |
| **Cluster size** | 112 | 86 | 144 |
| **Energy Weighted Score** | -947.3 | -1030.5 | -1092.6 |

| TRIM residue | LC3 residue | Contact Area | Exposed Area | % Buried | Distance(Å) |
|---|---|---|---|---|---|
| **N379** | N.D. | N.D. | N.D. | N.D. | N.D. |
| **Q380** | K55 | 40.63 | 101.7 | 40 | 2.457 |
| **W381** | D23 | 65.49 | 161 | 41 | 2.99 |
| **E382** | N.D. | N.D. | N.D. | N.D. | N.D. |
| **Q383** | N.D. | N.D. | N.D. | N.D. | N.D. |
| **L384** | R20 | 35.41 | 102.7 | 34 | 3.242 |

| B. Human TRIM47 (Alphafold: Q96LD4) x Human LC3B (PDB: 3VTU) | | | |
|---|---|---|---|
| **LIR motif position**: 379-384 **LIR sequence**: NQWEQL | | | |
| | **Balanced** | **Electrostatic** | **Hydrophobic** |
| **Cluster size** | 238 | 113 | 112 |
| **Energy Weighted Score** | -1301.7 | -966 | -1081.2 |

| TRIM residue | LC3 residue | Contact Area | Exposed Area | % Buried | Distance(Å) |
|---|---|---|---|---|---|
| **N379** | N.D. | N.D. | N.D. | N.D. | N.D. |
| **Q380** | N.D. | N.D. | N.D. | N.D. | N.D. |
| **W381** | I21 | 125.3 | 176.2 | 71 | 2.337 |
| **E382** | R72 | 18.55 | 87.68 | 21 | 2.961 |
| **Q383** | N.D. | N.D. | N.D. | N.D. | N.D. |
| **L384** | V93 | 12.12 | 128 | 9 | 4.073 |

| C. Mouse Trim47 (Alphafold: Q8C0E3) x Mouse LC3B (Alphafold: Q9CQV6) | | | |
|---|---|---|---|
| **LIR motif position**: 383-388 **LIR sequence**: SQWEQL | | | |
| | **Balanced** | **Electrostatic** | **Hydrophobic** |
| Cluster size | 169 | 147 | 179 |
| Energy Weighted Score | -946.7 | -1046.7 | -1368.9 |

| Trim residue | LC3 residue | Contact Area | Exposed Area | % Buried | Distance(Å) |
|---|---|---|---|---|---|
| **S383** | N.D. | N.D. | N.D. | N.D. | N.D. |
| **Q384** | N.D. | N.D. | N.D. | N.D. | N.D. |
| **W385** | F80 | 79.25 | 156.4 | 51 | 1.391 |
| **E386** | N.D. | N.D. | N.D. | N.D. | N.D. |
| **Q387** | N.D. | N.D. | N.D. | N.D. | N.D. |
| **L388** | M88 | 22.99 | 111.7 | 21 | 1.569 |

**Table S4. In silico docking simulation of protein-protein interaction between TRIM47 and LC3B of human and mouse**

*Above* the summary of protein-protein interaction cluster scores of (A, B) human TRIM47 x human LC3B and (C) mouse Trim47 x mouse LC3B based on ClusPro 2.0 simulation; *Below* the summary of residue contacts between TRIM47 and LC3B at the corresponding LIR motifs. As a positive control, human SQSTM (Alphafold: Q13501) x Human LC3B (PDB: 3VTU) interactions demonstrate a cluster size of 50, 97 and 78 and Energy Weighted Score of -931.6, -994.2 and -1319 for Balanced, Electrostatic-favored and Hydrophobic-favored mode, respectively.
